# Ventral Intermediate Nucleus structural connectivity-derived segmentation: anatomical reliability and variability

**DOI:** 10.1101/2021.05.02.442321

**Authors:** Salvatore Bertino, Gianpaolo Antonio Basile, Alessia Bramanti, Rossella Ciurleo, Adriana Tisano, Giuseppe Pio Anastasi, Demetrio Milardi, Alberto Cacciola

**Author notes:** **Correspondence** Alberto Cacciola, MD, Brain Mapping Lab, Department of Biomedical, Dental Sciences and Morphological and Functional, Images, University of Messina, Messina, Italy. These authors equally contributed to the present work (Salvatore Bertino, Gianpaolo Antonio Basile).

## Abstract

The Ventral intermediate nucleus (Vim) of thalamus is the most targeted structure for the treatment of drug-refractory tremors. Since methodological differences across existing studies are remarkable and no gold-standard pipeline is available, in this study, we tested different parcellation pipelines for tractography-derived putative Vim identification.

Thalamic parcellation was performed on a high quality, multi-shell dataset and a downsampled, clinical-like dataset using two different diffusion signal modeling techniques and two different voxel classification criteria, thus implementing a total of four parcellation pipelines. The most reliable pipeline in terms of inter-subject variability has been picked and parcels putatively corresponding to motor thalamic nuclei have been selected by calculating similarity with a histology-based mask of Vim. Then, spatial relations with optimal stimulation points for the treatment of essential tremor have been quantified. Finally, effect of data quality and parcellation pipelines on a volumetric index of connectivity clusters has been assessed.

We found that the pipeline characterized by higher-order signal modeling and threshold-based voxel classification criteria was the most reliable in terms of inter-subject reliability regardless data quality. The maps putatively corresponding to Vim were those derived by precentral- and dentate nucleus-thalamic connectivity. However, tractography-derived functional targets showed remarkable differences in shape and sizes when compared to a ground truth model based on histochemical staining on seriate sections of human brain. Thalamic voxels connected to contralateral dentate nucleus resulted to be the closest to literature-derived stimulation points for essential tremor but at the same time showing the most remarkable inter-subject variability. Finally, the volume of connectivity parcels resulted to be significantly influenced by data quality and parcellation pipelines. Hence, caution is warranted when performing thalamic connectivity-based segmentation for stereotacting targeting.

## Introduction

Functional neurosurgery techniques, such as deep brain stimulation (DBS) and magnetic resonance - guided focused ultrasounds (MRgFUS), have been increasingly used as effective therapeutic strategies for treating severe, drug-refractory tremors (Benabid et al., 1996; Elias et al., 2013). Given its central role in tremor circuitry, the Ventral intermediate nucleus (Vim) of the thalamus is the most commonly targeted structure for treatment of essential tremor (ET) and tremor-dominant Parkinson’s disease (PD) (Cury et al., 2017). The Vim, as described in Hassler’s classification, belongs to motor thalamic nuclei and it is depicted as a wedge-shaped area, located at the inferior edge of ventrolateral thalamus (Hassler, 1983). Classical tract-tracing and immunohistochemical studies, conducted both on non-human and human primates, define the Vim as the entry zone of the dento-rubro-thalamo-cortical (DRTC) tract (Calzavara et al., 2005; Darian-Smith et al., 1990; Dum and Strick, 2003; Gallay et al., 2008).

As conventional magnetic resonance imaging (MRI) lacks of intrinsic contrast to identify thalamic nuclei, interest towards advanced MRI techniques for personalized targeting is currently growing (Boutet et al., 2019; Fenoy and Schiess, 2018; Shah et al., 2020). Among these, structural connectivity-based-parcellation (CBP), which relies on the classification of thalamic voxels according to their connectivity profile, allows the identification of a discrete number of connectivity parcels whose spatial relation could be comprehensively investigated (Cacciola et al., 2019b; da Silva et al., 2017; Middlebrooks et al., 2018b; Plantinga et al., 2018; Pouratian et al., 2011). While many works have applied structural connectivity-based methods to identify the Vim in a clinical research context, a well-established and clinically suitable pipeline for thalamic parcellation is still lacking as significant methodological heterogeneity is present across different works (Akram et al., 2019; Krishna et al., 2019; Sammartino et al., 2016; Tian et al., 2018). Many methodological variables are known to have a strong impact on tractography-based parcellation results, such as acquisition parameters (b-value, spatial and angular resolution) (Ambrosen et al., 2020; Essayed et al., 2017), diffusion signal modelling (Behrens et al., 2007; Jbabdi and Johansen-Berg, 2011), tractography algorithm (Girard et al., 2020; Petersen et al., 2017) and voxel classification criteria (Bertino et al., 2020; Jbabdi and Johansen-Berg, 2011; Jeurissen et al., 2019; Plantinga et al., 2018). However, studies investigating systematically the impact of such variables on the results of the output thalamic parcellations are currently lacking. To serve as a good proxy for Vim identification, a “candidate” pipeline should rely on a combination of such variables resulting in connectivity maps that are reliable across subjects, adherent to the anatomical ground-truth and as close as possible to clinically effective stimulation/ablation point. In addition, it should provide interpretable results and be also sufficiently easy to perform in advanced-care clinical contexts. Indeed, while most of the existing works employed high angular resolution diffusion imaging datasets (HARDI) (Jones et al., 1999) with high-order diffusion modelling techniques (Akram et al., 2019; Behrens et al., 2007; Middlebrooks et al., 2018b; Schlaier et al., 2017; Tournier et al., 2007), lower quality datasets and diffusion tensor modelling are still employed, as they require less computational power, are faster to implement, and, in general, more available in clinical settings (Fenoy and Schiess, 2017; Miller et al., 2019). A comparison of lower-level methods versus state-of-art tractography for thalamic parcellation may help in translating connectivity-derived thalamic parcellation into a reliable clinical routine.

Herein, we tested four different parcellation pipelines for tractography-aided identification of thalamic functional targets to disentangle the effects of acquisition parameters, diffusion signal modelling and voxel classification criteria on parcellation results. Specifically, we evaluated different parcellation protocols in terms of between-subjects similarity, with the aim of identifying a reliable protocol able to work both on high quality and “clinical-like” data. In particular, we tested a hypothesis-driven thalamic parcellation method (Behrens et al., 2003) on a high quality, multi-shell dataset and a downsampled, clinical-like dataset of 210 healthy subjects of the Human Connectome Project (HCP) repository combining two different signal modelling techniques and two different voxel classification criteria. CBP pipelines were tested for reliability, and the best working approach in terms of inter-subject similarity was picked for further analyses such as similarity with histology-based Vim, group-level versus individualized maps similarity and spatial relations with literature-based optimal stimulation points.

## Materials and Methods

### Subjects

We employed diffusion and structural MRI data of 210 healthy subjects (males = 92, females = 118, age range 22-36 years) obtained from the HCP repository. Data have been acquired by the Washington University, University of Minnesota and Oxford university (WU-Minn) HCP consortium. Subject recruitment procedures, informed consent and sharing of de-identified data were approved by the Washington University in St. Louis Institutional Review Board (IRB) (Van Essen et al., 2013).

### High quality HCP data

MRI acquisitions were carried out using a custom-made Siemens 3T “Connectome Skyra” (Siemens, Erlangen, Germany), provided with a Siemens SC72 gradient coil and stronger gradient power supply with maximum gradient amplitude (Gmax) of 100 mT/m (initially 70 mT/m and 84 mT/m in the pilot phase), which allows improvement of diffusion-weighted imaging (DWI).

DWIs were acquired using a single-shot 2D spin-echo multiband Echo Planar Imaging (EPI) sequence and equally distributed over 3 shells (*b-values* 1000, 2000, 3000 mm/s^2^), 90 directions per shell, spatial isotropic resolution of 1.25 mm (Sotiropoulos et al., 2013).

High resolution T1-weighted MPRAGE images were collected using the subsequent parameters: voxel size = 0.7 mm, TE = 2.14 ms, TR = 2,400 ms (Van Essen et al., 2012).

Data were acquired in the minimally preprocessed form, briefly, distortion correction, motion correction, brain extraction and registration of structural and diffusion data to one another and to MNI space were already carried out in the downloaded package (Glasser et al., 2016, 2013; Sotiropoulos et al., 2013; Van Essen et al., 2012).

### Structural and diffusion data post-processing

T1-weighted structural images underwent cortical and subcortical segmentation implemented by FAST and FIRST FSL functions respectively (https://fsl.fmrib.ox.ac.uk/fsl) (Patenaude et al., 2011; Smith, 2002; Smith et al., 2004). T1-weighted volumes were also coregistered to the ICBM 2009b nonlinear asymmetric template (Fonov et al., 2009) using symmetric diffeomorphic image registration (SyN) normalization (Avants et al., 2009) and direct and inverse transformations were saved for further use. Thalamic regions of interest (ROIs) were retrieved from FSL FIRST tool, which allows, for each subject, the segmentation of fifteen subcortical structures in a fully automated fashion (Patenaude et al., 2011). Details about the other ROIs employed in this work can be found in Supplementary Information.

Two different signal modeling algorithms were applied on diffusion datasets: a traditional, diffusion-tensor based model (DTI) (Basser et al., 1994) and a high-order model based on multi-shell, multi tissue constrained spherical deconvolution (MSMT-CSD). CSD signal modeling estimates white matter fibre Orientation Distribution Function (fODF) from the diffusion-weighted deconvolution signal using a single fiber response function (RF) as reference (Tournier, Calamante, & Connelly, 2007; Tournier et al., 2008). CSD signal modelling allows to reconstruct successfully complex fibers configurations, especially crossing-fibers (Farquharson et al., 2013). MSMT-CSD represents a further development of the conventional CSD approach calculating different response functions for gray matter (GM), white matter (WM) and cerebrospinal fluid (CSF) allowing for more precise fODF orientation reducing the presence of spurious fODF in voxels which do not contain white matter (Jeurissen et al., 2014). Both tensor fitting, response function calculation, fODF estimation and tractography were implemented using MRtrix3 (www.mrtrix.org).

### Downsampled HCP data

In order to test how different pipelines would perform on lower quality datasets, diffusion data were downsampled to imitate the most relevant features of low-resolution, non-HARDI scans which are commonly available in advanced-care clinical settings. To this purpose, a single DWI shell (b= 1000 s/mm^2^) has been extracted including six b0 volumes, spatial resolution was lowered by reducing voxel size to 2.00 mm isotropic and angular resolution was decreased retaining only 26 unique diffusion weighted volumes from the selected shell (Wasserthal et al., 2018). Since white matter modeling of traditional CSD may be affected in areas characterized by partial volume effects, and MSMT-CSD requires multi-shell data which are rarely acquired during clinical routine, we modeled diffusion signal using single shell 3-tissue CSD (SS3T-CSD). In such variant of the CSD-method, response functions for single-fibre WM as well as GM and CSF are estimated from the data using an unsupervised method (Dhollander et al., 2019). SS3T-CSD signal modeling was performed using MRtrix3Tissue (https://3Tissue.github.io), a fork of MRtrix3 (Tournier et al., 2019).

### Tractography

Seed-based probabilistic tractography has been employed to reconstruct streamlines joining the thalamus to cortical targets and to the contralateral superior cerebellar peduncle (SCP) ROI. Tractography has been performed with default tracking parameters using default probabilistic algorithms available in the MrTrix3 software (iFOD2 for CSD data; Tensor-prob for tensor-based tractography). Both in CSD- and DTI-based approaches, streamlines joining thalamus to cortical areas have been extracted using default parameters, seeding from thalamus and selecting a given cortical area as an inclusion region. On the other hand, when CSD signal modeling was implemented, DRTC tract has been reconstructed seeding from SCP ROIs (Palesi et al., 2016, 2015), using masks of the contralateral red nucleus and thalamus as inclusion regions; when tensor model was fitted, streamlines were seeded throughout the SCP mask, using the ipsilateral red nucleus and thalamus as inclusion regions. This choice was motivated by the inability of tensor-based tractography in resolving crossing fibers and has been already implemented in previous studies employing DTI signal modeling (Krishna et al., 2019; Palesi et al., 2015; Sammartino et al., 2016).

### Connectivity-based parcellation

Hypothesis-driven CBP of thalamus was performed according to its structural connectivity to eight cortical targets and to SCP, thus, for each subject, nine connectivity parcels were obtained.

CBP has been performed on each subject, applying two distinct signal modeling techniques (CSD, DTI). In addition, each pipeline was carried out using two voxel classification criteria: threshold-based approach and hard-segmentation. The former simply rules out voxels characterized by low connectivity values by applying an arbitrary threshold, that was set to 25%, as in previous works (Patriat et al., 2018; Plantinga et al., 2018). The hard segmentation approach uses a winner-takes-all strategy implemented on FSL to attribute voxels univocally to targets with the highest track-density (Behrens et al., 2003).

In summary, a total of four pipelines were implemented for each subject both on high-quality and on clinical-like datasets. For the sake of clarity, from now on, we will refer to the tested pipelines as follows:

- CSD-THR
- CSD-WTA
- DTI-THR
- DTI-WTA

Where “THR” refers to the threshold-based approach and “WTA” stands for “winner-takes-all” i.e. the hard-segmentation strategy.

The parcellation procedure was implemented as summarized below:

1. Streamlines connecting thalamus to ipsilateral cortical areas were reconstructed using seed-based tractography; in particular, thalamus was set as a seed region, whilst a selected cortical target was set as an include region; this process was re-iterated for each cortical target. DRTC was reconstructed using SCP as a seed region and contralateral red nucleus and thalamus as inclusion regions. Contralateral thalamus was used as exclusion region. Ipsilateral red nucleus and thalamus were used as inclusion regions in DTI-based pipelines.
2. The obtained streamlines were used as a form of contrast to obtain a track density weighted map for each pathway of interest (Calamante et al., 2010).
3. Track density maps were multiplied for thalamus masks using *fslmaths* command, thus allowing the retrieval of thalamic track density-weighted maps.
4. TDI maps were normalized by calculating for each map the ratio between its voxels’ intensity and its mean intensity to mitigate overestimation/underestimation effects due to targets volumes thus obtaining comparable values among maps.
5. In -THR pipelines, the normalized maps underwent a threshold of 25% to remove voxels characterized by lower track density, whilst in -WTA the find-the-biggest algorithm was applied to attribute voxels to the connectivity clusters characterized by highest number of streamlines.
6. All connectivity maps derived from the previous step were binarized and registered to the ICBM 2009b nonlinear asymmetric template (Fonov et al., 2009) using nonlinear transformations obtained from structural registration.
7. Once registered to standard space, binarized maps were summed up to obtain group-level probability maps. Finally, a 50% threshold has been applied to such maps to retain only voxels overlapping in at least 50% of the sample thus obtaining maximum probability maps (MPMs) (da Silva et al., 2017; Domin and Lotze, 2019). Such MPMs served as reference group-level maps for further analysis (see atlas-based and personalized procedures). Finally, volumes and center of gravity (COG) of MPMs, for each pipeline have been quantified.

### Reliability analysis

Reliability across parcellation pipelines and connectivity clusters has been evaluated both in high quality and downsampled datasets, by calculating inter-subject similarity metrics derived from Tanimoto coefficient (Crum et al., 2006). Such analysis has been conducted after co-registration of the subject-level connectivity clusters to ICBM template, i.e. maps obtained in the abovementioned step 6 of CBP.

At first, “parcellation-level”, groupwise overlap was measured calculating the total accumulated overlap (TAO) for each pipeline. For a group of *m* pairs of images, where *m* represents all the possible pairwise combinations between images of the same cluster, TAO is defined as:

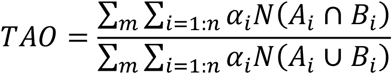

where *n* is the total number of clusters obtained from a given parcellation pipeline (CSD-THR, CSD-WTA, DTI-THR, DTI-WTA), *i* is the cluster label and α is a weighting coefficient. We defined α as the inverse of the mean of the absolute value of volumes for A and B, to avoid overestimation of larger parcels:

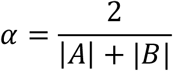

The reliability of each connectivity parcel across all parcellation pipelines tested has been assessed by calculating overlap by label (OBL) which is defined as:

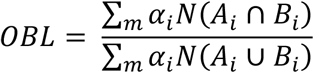

For both measures, the result is a number ranging between 0 and 1, where a value closer to 1 indicates higher similarity whilst a value closer to 0 indicates high dissimilarity (Bertino et al., 2020; da Silva et al., 2017).

### Similarity between connectivity patterns and histologically-defined Vim

Among thalamic connectivity maps, we identified those which showed higher spatial similarity to Vim thalamic nucleus and could be then more suitable as a connectivity-derived proxy for Vim identification. Spatial similarity was evaluated by comparing individual connectivity maps and a publicly available Vim ROI obtained by digitizing high-resolution thin-slice histological data from a single post mortem specimen (Chakravarty et al., 2006) and made available as part of a standard-space MRI atlas (Ewert et al., 2018). Inverse transformations obtained from *SyN* registration have been employed to register the Vim ROI of each hemisphere as provided by the DISTAL atlas from 2009b MNI non-linear template to the native space of each subject. Finally, similarity of the Vim masks with individualized connectivity maps obtained after the normalization step has been assessed calculating average Dice similarity coefficients (Dice, 1945):

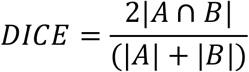

Spatial maps showing higher similarity to histological Vim (average Dice coefficient ≥ 0.2 bilaterally and both in high-quality and downsampled data) were picked up for further analysis.

### Variability across individualized connectivity maps

To evaluate differences between group-level and individualized thalamic parcels, i.e. how well the group-level MPMs were representative of subject-level connectivity maps we sought to assess spatial similarity between group level and individualized maps. In order to achieve this, the inverse transformations obtained from *SyN* have been used to register MPMs derived from the most reliable approach to each subject native space, then, similarity between atlas-based (MPMs) and individualized connectivity maps has been assessed by calculating Dice coefficient. In addition, the COGs of connectivity maps registered on the ICBM template have been extracted for each subject and pairwise Euclidean distances were also calculated between individual COGs and the COGs extracted from maximum probability maps.

To further evaluate between-subject variability in position of COGs, the mean displacement of individual COGs from average group-level coordinates was also calculated. Group level coordinates were obtained by averaging the individual COGs across all subjects. Hence, pairwise Euclidean distances between group level-COGs and COGs of each individual connectivity cluster have been assessed at subject level and then averaged.

### Spatial relations between MPMs and optimal stimulation points

In order to investigate if the location of group-level connectivity maps match the stimulation points for essential tremor (ET) available in literature, we calculated Euclidean distances between MPM COGs and three sets of coordinates provided by a recent study (Elias et al., 2020). Specifically, the first set of coordinates was represented by the “classic” literature-based optimal lead location, obtained from 37 patients affected by ET and translated to MNI space using a probabilistic conversion method (Horn et al., 2017; Papavassiliou et al., 2004). The second and third sets of coordinates were derived from the application of volume of tissue activated (VTA) probabilistic stimulation mapping on a cohort of 39 ET patients: the former was derived from an unweighted frequency map and the latter from clinically weighted hotspots. While the unweighted frequency maps represent the most frequently stimulated voxels across the cohort, the clinically weighted hotspots represents the voxels whose stimulation led to above-mean clinical improvement (Elias et al., 2020).

### Effect of pipelines and data quality on connectivity map volumes

In order to obtain quantitative estimates of connectivity map volumes, streamline density index (SDI), which consists of the percentage volume ratio between connectivity maps and thalamus ROI, was calculated at subject level for all pipelines, both in high quality and downsampled datasets using the formula below (Theisen et al., 2017):

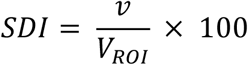

Where *v* is the volume (in voxels) of the parcel after threshold application or hard segmentation, namely, clusters obtained from the step 5 of the parcellation pipelines (see above); V_ROI_ is the volume (in voxels) of the thalamus ROI. A two-way repeated measures ANOVA was conducted to examine the effect of parcellation pipelines and data quality on SDI. For the selected connectivity clusters of each hemisphere a distinct general linear model was computed, setting SDI as dependent variable and data quality and pipeline as within-subject factors with two (high quality and downsampled) and four (CSD-THR, CSD-WTA, DTI-THR, DTI-WTA) levels respectively. A p-value of <0.05 was deemed to be significant and, if necessary, when pairwise comparisons were performed on pipelines, p-values were adjusted for multiple comparisons using Bonferroni correction. Statistical analysis has been carried out using SPSS Statistics (IBM SPSS Statistics for Windows, Version 25.0. Armonk, NY: IBM Corp).

## Results

### Thalamic connectivity-based parcellation

The full, stepwise, workflow adopted for both high quality and downsampled datasets is summarized in Figure 1.

**Figure 1.**
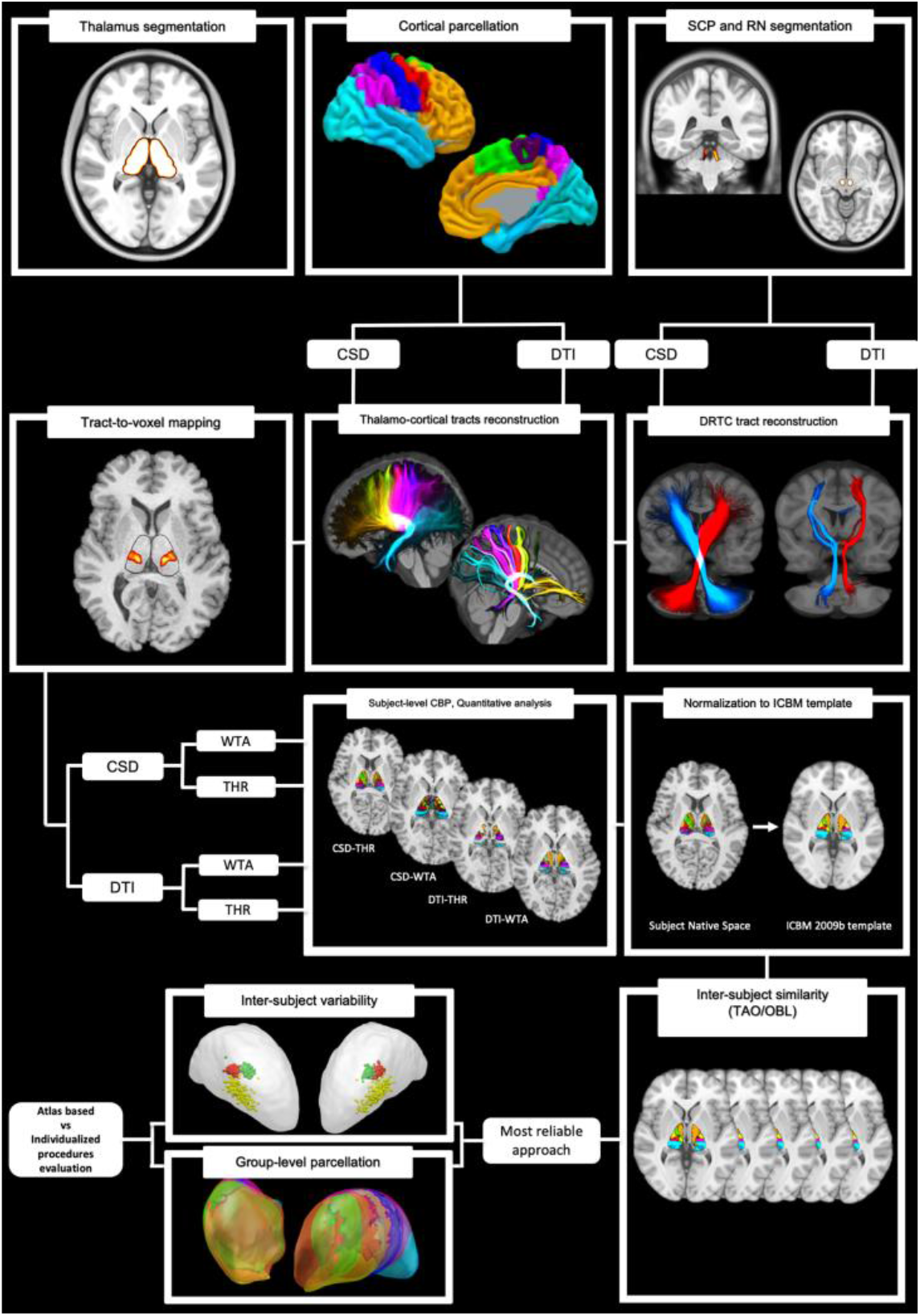
Experimental workflow. Once regions of interest were selected, tractographic reconstruction of cortico-thalamic and dento-rubro-thalamo-cortical pathways was performed using two different signal modeling techniques (CSD/DTI). Then, streamlines connecting thalamus to each region of interest were used as a contrast to obtain track-density weighted maps. For each obtained connectivity map, two different voxel classification criteria were applied (THR, WTA) thus providing four distinct parcellation solutions for each subject. The resulting maps were employed to evaluate quantitative differences across different pipelines and data quality. All the obtained maps were registered to ICBM 2009b template and binarized; once registered on the common standard space, such clusters underwent reliability analysis. Maps derived from most reliable approach underwent inter-subject variability analysis; then, have been summed up to obtain maximum probability maps (MPM). These group-level maps have been further employed to investigate atlas-based vs individualized differences.

Each parcellation pipeline brought nine connectivity clusters, each one exhibiting peculiar shape, dimensions and orientation along the anterior-posterior thalamic axis. The whole parcellation for the most reliable approach (see next paragraph) is shown in Figure 2 while spatial maps of each connectivity parcel are provided in Supplementary Figures 1-2. Regardless data quality, the spatial organization of connectivity maps followed, at least in part, the antero-posterior arrangement of cortical targets; besides, all parcels extended both in dorsal and ventral thalamic aspects, except for the dentate MPMs which occupied the most ventral and lateral thalamic portions. Overall, the parcellation here obtained shared similarities with those provided by existing studies (Behrens et al., 2003; Broser et al., 2011; Middlebrooks et al., 2018a; Traynor et al., 2010).

**Figure 2.**
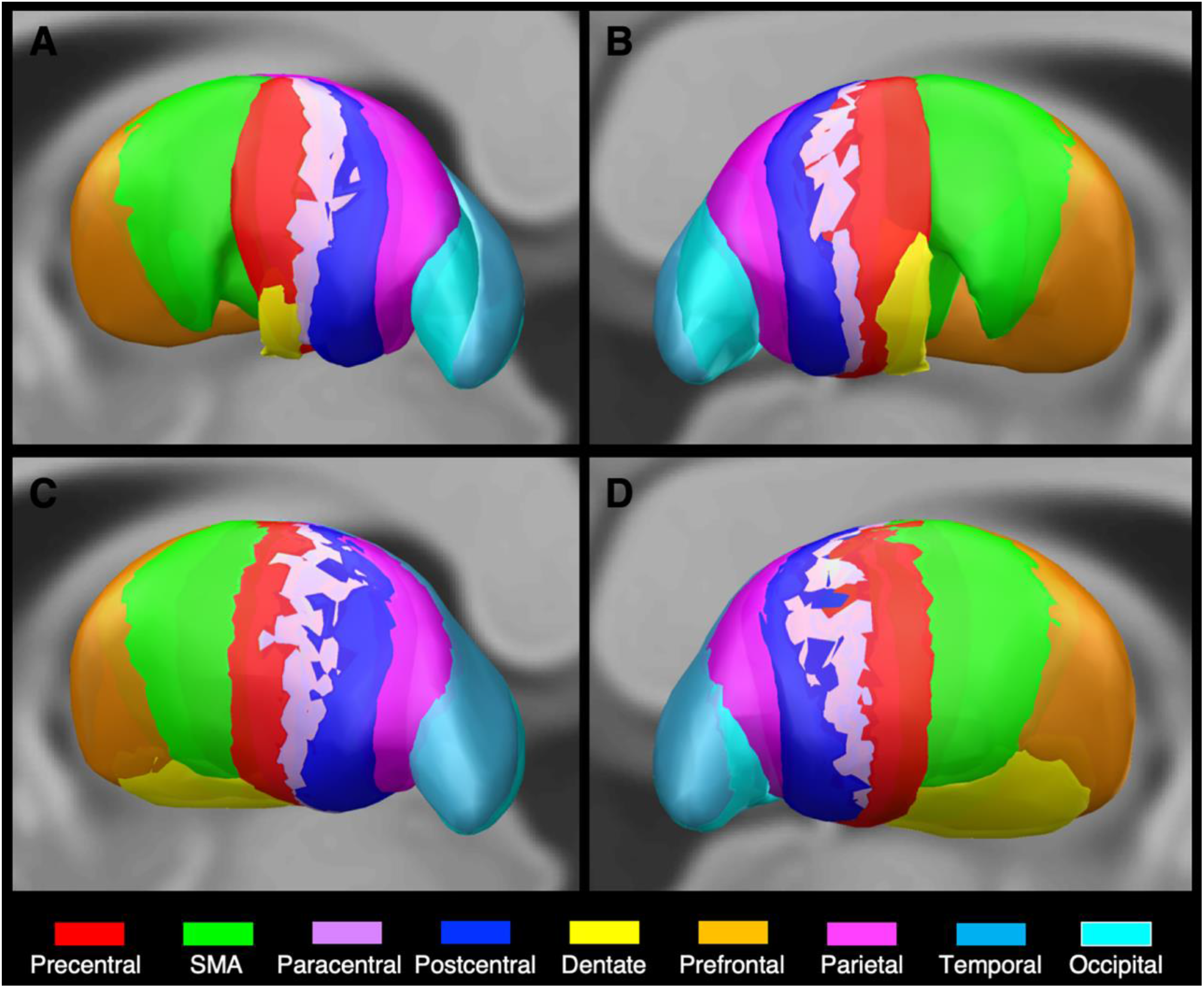
Group level thalamic CBP. 3D reconstruction of MPMs obtained from the most reliable approach (CSD-THR). The upper row depicts mid-sagittal sections of left (A) and right (B) MPMs obtained from high quality datasets. The bottom row shows mid-sagittal sections of left (C) and right (D) MPMs retrieved from downsampled data. MPMs have been labeled according to the following color code: precentral (red), SMA (green), paracentral (pink), postcentral (blue), dentate (yellow), prefrontal (orange), parietal (purple), temporal (light blue), occipital (cyan).

### Parcellation-level and cluster-level reliability of parcellation

Once thalamic CBP has been performed on each subject of both datasets, TAO has been calculated, returning a number between 0 and 1 for each of the pipelines tested (Figure 3A, Table 1).

**Table 1.** The table shows the values of total accumulated overlap (TAO) across different parcellation pipelines (CSD-THR, CSD-WTA, DTI-THR, DTI-WTA) and datasates (high quality and downsampled).

| Total<br>accumulated<br>overlap<br>(TAO) | High Quality |  | Downsampled |  |
| --- | --- | --- | --- | --- |
|  | Left | Right | Left | Right |
| **CSD-THR** | 0.37 | 0.36 | 0.39 | 0.39 |
| **CSD-WTA** | 0.29 | 0.29 | 0.17 | 0.18 |
| **DTI-THR** | 0.13 | 0.13 | 0.12 | 0.12 |
| **DTI-WTA** | 0.19 | 0.19 | 0.18 | 0.18 |

**Figure 3.**
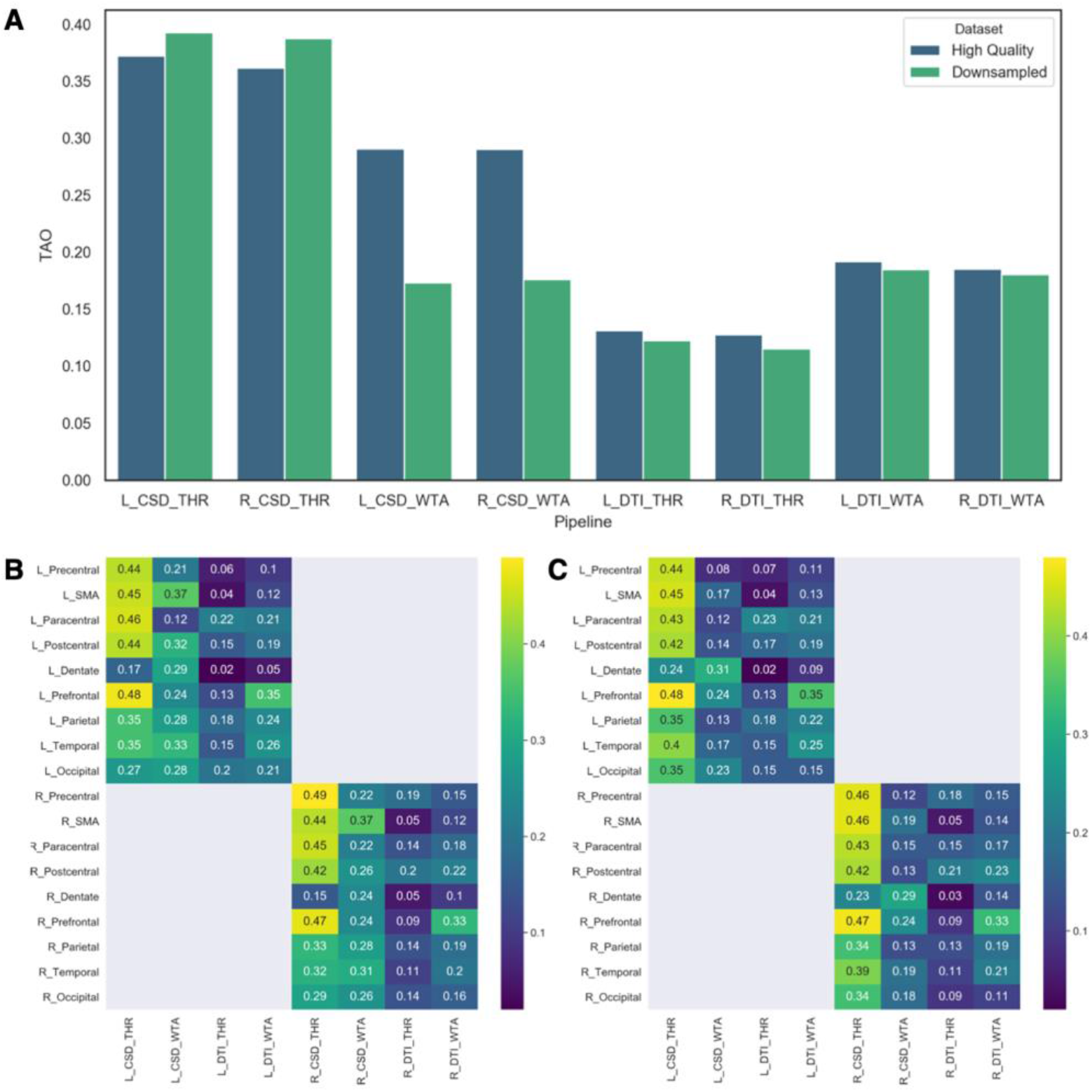
Parcellation-level and cluster-level reliability of parcellation. (A) Bar plots showing values for parcellation-wise, total accumulated overlap (TAO) for each parcellation pipeline, both in high quality (Blue) and downsampled (Green) datasets are provided. In the bottom part of the figure, heatmaps reporting values of cluster-wise overlap-by-label (OBL) for high quality (B) and downsampled (C) datasets are supplied. As shown by the color bar, dark blue cells display clusters with the lowest degree of inter-subject similarity, which, as increases, becomes yellow in cells corresponding to clusters exhibiting the highest inter-subject similarity.

Across high quality datasets, TAO ranged from 0.13 to 0.37, with CSD-based pipelines systematically exhibiting higher values than DTI-based ones. In particular, the highest TAO was reached by CSD-THR pipelines (Left 0.37, Right 0.36), followed by CSD-WTA pipelines (Left 0.29, Right 0.29). Overall, DTI-based pipelines exhibited lower TAO values, with WTA approaches being slightly higher (Left 0.19, Right 0.19) than those employing the adaptive threshold (Left 0.13, Right 0.13). Concerning downsampled datasets, TAO values were lower than those displayed in high quality datasets, except for the CSD-THR pipeline which showed slightly higher values than those obtained for high quality datasets (Left 0.39, Right 0.39). The CSD-WTA pipeline showed the most remarkable difference between high quality (Left 0.29, Right 0.29) and downsampled data (Left 0.17, Right 0.18). On the other hand, although being lower, the difference in TAO values with high quality datasets in DTI-THR (Left 0.12, Right 0.12) and DTI-WTA (Left 0.18, Right 0.18), pipelines is minimal (Figure 3A).

Summarizing, the highest TAO values have been observed for both kind of datasets with CSD-THR; thus, connectivity maps obtained using such combination have been considered for further analyses (see following paragraphs).

Cluster-level reliability has been assessed by calculating OBL (Figure 3B-C, Table 2). Such inter-subject similarity metric exhibited values ranging from 0.05 to 0.49 for high quality datasets and from 0.02 to 0.48 for downsampled datasets. The pattern described for TAO, with higher scores achieved by CSD-THR pipelines both in high quality and downsampled data, was maintained in all connectivity clusters considered, with the only exception of dentate connectivity maps. Indeed, while other motor-related connectivity maps displayed fair inter-subject similarity, in CSD-THR approaches, the dentate maps showed overall low OBL, with the highest values displayed by CSD-WTA pipelines both in high quality (Left 0.29, Right 0.24) and downsampled datasets (Left 0.31, Right 0.29).

**Table 2.**
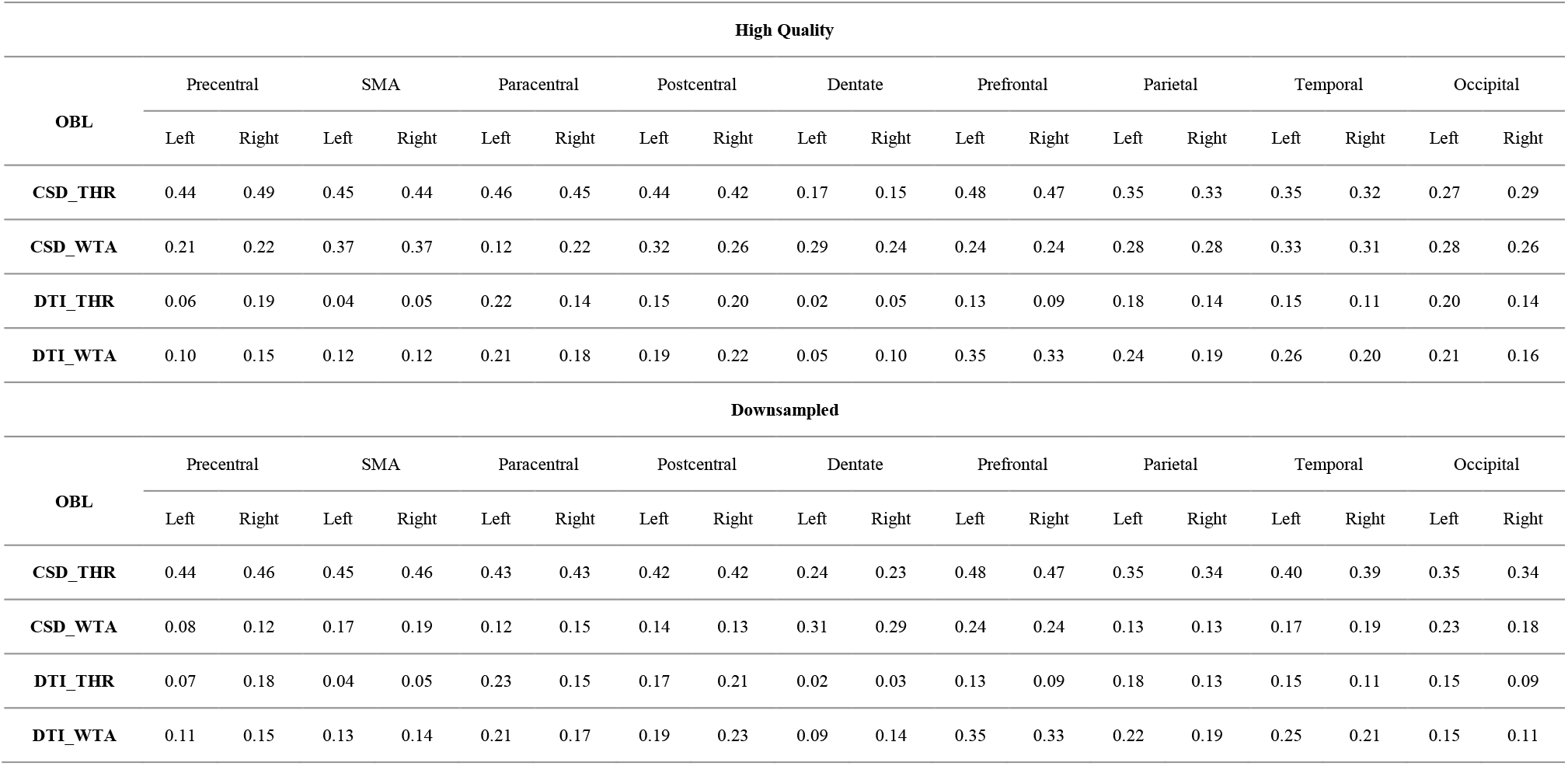
Table showing values of overlap by label (OBL) across different clusters for both high quality and downsampled datasets.

| High Quality |  |  |  |  |  |  |  |  |  |  |  |  |  |  |  |  |  |  |
| --- | --- | --- | --- | --- | --- | --- | --- | --- | --- | --- | --- | --- | --- | --- | --- | --- | --- | --- |
| OBL | Precentral |  | SMA |  | Paracentral |  | Postcentral |  | Dentate |  | Prefrontal |  | Parietal |  | Temporal |  | Occipital |  |
|  | Left | Right | Left | Right | Left | Right | Left | Right | Left | Right | Left | Right | Left | Right | Left | Right | Left | Right |
| CSD_THR | 0.44 | 0.49 | 0.45 | 0.44 | 0.46 | 0.45 | 0.44 | 0.42 | 0.17 | 0.15 | 0.48 | 0.47 | 0.35 | 0.33 | 0.35 | 0.32 | 0.27 | 0.29 |
| CSD_WTA | 0.21 | 0.22 | 0.37 | 0.37 | 0.12 | 0.22 | 0.32 | 0.26 | 0.29 | 0.24 | 0.24 | 0.24 | 0.28 | 0.28 | 0.33 | 0.31 | 0.28 | 0.26 |
| DTI_THR | 0.06 | 0.19 | 0.04 | 0.05 | 0.22 | 0.14 | 0.15 | 0.20 | 0.02 | 0.05 | 0.13 | 0.09 | 0.18 | 0.14 | 0.15 | 0.11 | 0.20 | 0.14 |
| DTI_WTA | 0.10 | 0.15 | 0.12 | 0.12 | 0.21 | 0.18 | 0.19 | 0.22 | 0.05 | 0.10 | 0.35 | 0.33 | 0.24 | 0.19 | 0.26 | 0.20 | 0.21 | 0.16 |
| Downsampled |  |  |  |  |  |  |  |  |  |  |  |  |  |  |  |  |  |  |
| OBL | Precentral |  | SMA |  | Paracentral |  | Postcentral |  | Dentate |  | Prefrontal |  | Parietal |  | Temporal |  | Occipital |  |
|  | Left | Right | Left | Right | Left | Right | Left | Right | Left | Right | Left | Right | Left | Right | Left | Right | Left | Right |
| CSD_THR | 0.44 | 0.46 | 0.45 | 0.46 | 0.43 | 0.43 | 0.42 | 0.42 | 0.24 | 0.23 | 0.48 | 0.47 | 0.35 | 0.34 | 0.40 | 0.39 | 0.35 | 0.34 |
| CSD_WTA | 0.08 | 0.12 | 0.17 | 0.19 | 0.12 | 0.15 | 0.14 | 0.13 | 0.31 | 0.29 | 0.24 | 0.24 | 0.13 | 0.13 | 0.17 | 0.19 | 0.23 | 0.18 |
| DTI_THR | 0.07 | 0.18 | 0.04 | 0.05 | 0.23 | 0.15 | 0.17 | 0.21 | 0.02 | 0.03 | 0.13 | 0.09 | 0.18 | 0.13 | 0.15 | 0.11 | 0.15 | 0.09 |
| DTI_WTA | 0.11 | 0.15 | 0.13 | 0.14 | 0.21 | 0.17 | 0.19 | 0.23 | 0.09 | 0.14 | 0.35 | 0.33 | 0.22 | 0.19 | 0.25 | 0.21 | 0.15 | 0.11 |

### Connectivity-derived Vim identification

To identify connectivity parcels that could best suit as a proxy for Vim identification, we calculated average Dice coefficients between subject-level connectivity parcels (obtained using CSD-THR pipeline) and histology-based ROIs of Vim registered on each subject’s native space (Table 3).

**Table 3.** Average Dice coefficients between histology-based Vim registered to each subject native space and each connectivity parcels for the most reliable pipeline (CSD-THR) in high quality and downsampled datasets.

| AvgDICE | High Quality |  | Downsampled |  |
| --- | --- | --- | --- | --- |
| Clusters | Left | Right | Left | Right |
| Precentral | 0.25 | 0.30 | 0.27 | 0.31 |
| SMA | 0.22 | 0.20 | 0.26 | 0.27 |
| Paracentral | 0.18 | 0.20 | 0.24 | 0.27 |
| Postcentral | 0.08 | 0.11 | 0.20 | 0.23 |
| Dentate | 0.27 | 0.27 | 0.25 | 0.32 |
| Prefrontal | 0.14 | 0.12 | 0.17 | 0.16 |
| Parietal | 0.01 | 0.00 | 0.08 | 0.07 |
| Temporal | 0.00 | 0.00 | 0.04 | 0.03 |
| Occipital | 0.00 | 0.00 | 0.04 | 0.04 |

Overall, connectivity-derived maps exhibited low similarity with Vim, with Dice values ranging from 0 to 0.30 for high quality dataset and from 0.03 to 0.32 for downsampled data. Among connectivity maps, the highest Dice values (≥ 0.2 bilaterally and both in high-quality and downsampled data) were obtained by the precentral, dentate, and SMA clusters (Figure 4); these parcels were subsequently picked up for further analysis.

**Figure 4.**
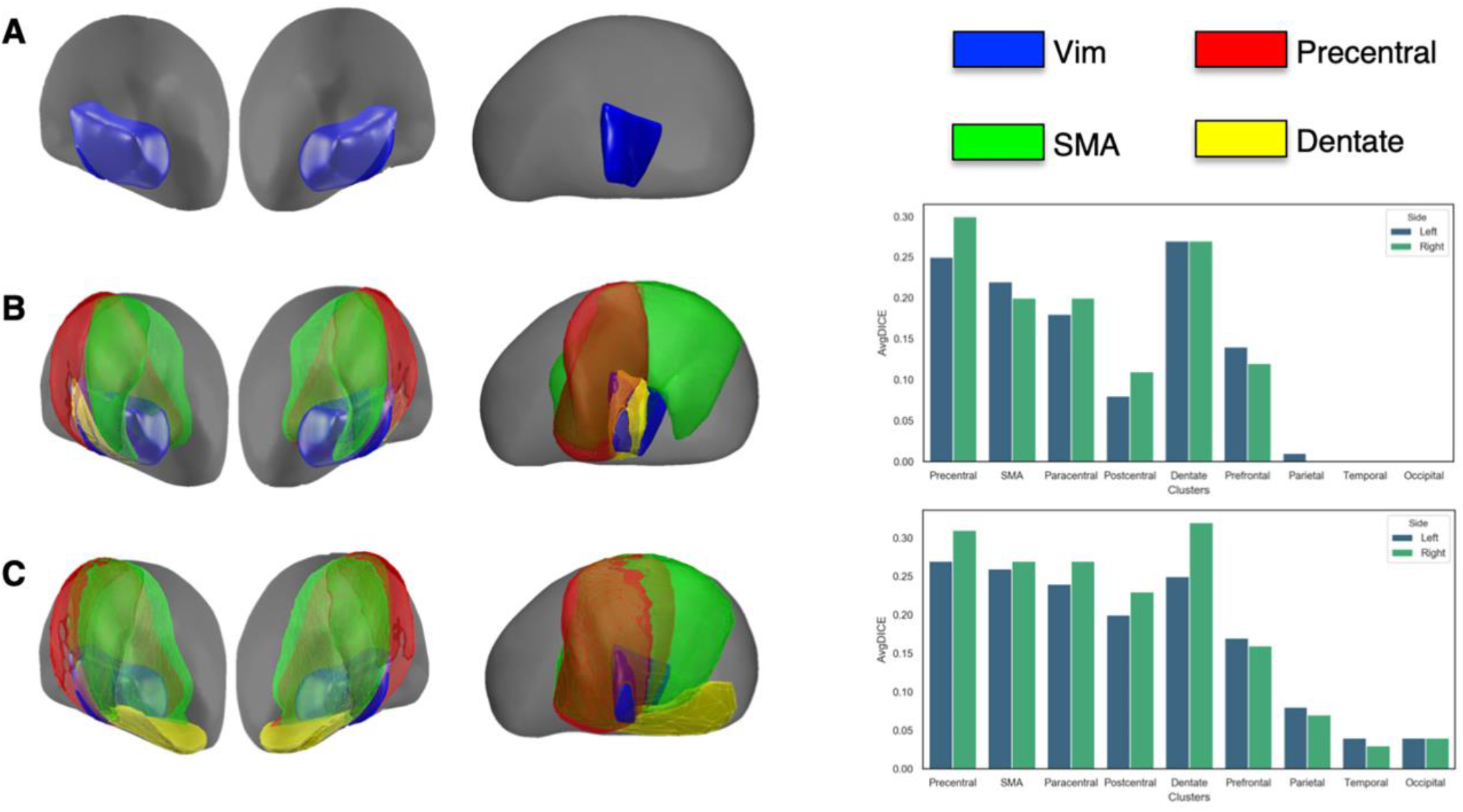
Comparison between histologic atlas-based and personalized procedures. This figure depicts spatial relations between a histology-based map of the Vim and the main “tremor-related” connectivity maps. For visualization purposes, the Vim has been superimposed on ICBM 2009b asymmetric template with precentral (red), SMA (green) and dentate (yellow) MPM derived from the most reliable pipeline. The upper row shows the Vim as included in the DISTAL atlas (A). The middle row shows frontal and lateral views of thalamus with Vim and MPMs obtained from high quality data superimposed (B). The lower row shows frontal and lateral views of thalamus with Vim and MPMs obtained from downsampled data (C). On the extreme right, bar plots summarizing average Dice coefficient between Vim and individualized clusters derived from high quality (B) and downsampled (C) are provided.

Notice that, amongst the remaining parcels, lesser degree of similarity had been observed for prefrontal and paracentral maps (≥ 0.2 in downsampled, but not in high-quality data) that were therefore not considered for further analysis, whilst minimal to none overlap was demonstrated between Vim and thalamic territories connected to postcentral, parietal, temporal and occipital cortices. Volumes and COG of the group-level putative Vim MPMs are summarized in Table 4 whilst those of the remaining MPMs are reported in Supplementary Table 1.

**Table 4.**
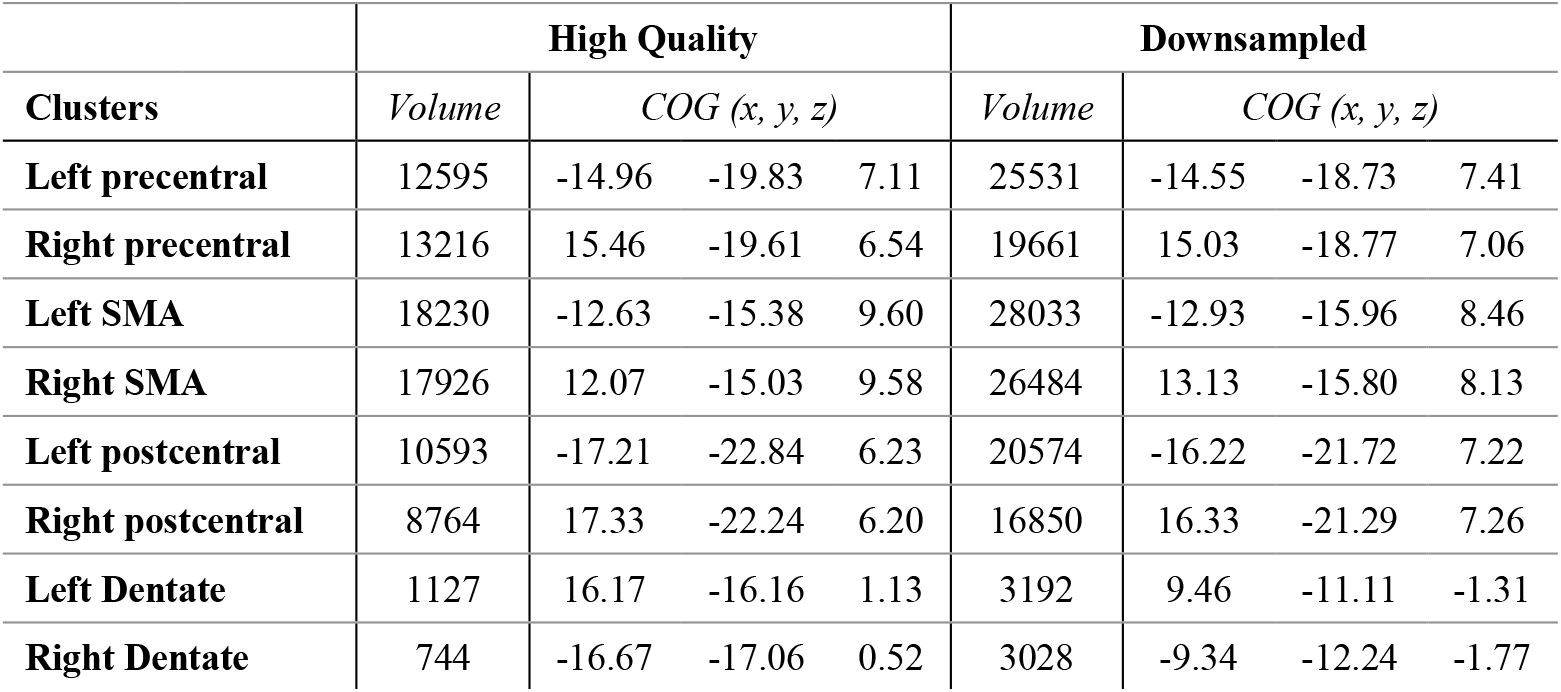
Volumes (in voxels) and COG (x, y, z) of sensorimotor MPMs obtained from the most reliable pipeline (CSD-THR), for high quality and downsampled dataset.

|  | High Quality |  |  |  | Downsampled |  |  |  |
| --- | --- | --- | --- | --- | --- | --- | --- | --- |
| Clusters | Volume | COG (x, y, z) |  |  | Volume | COG (x, y, z) |  |  |
| Left precentral | 12595 | -14.96 | -19.83 | 7.11 | 25531 | -14.55 | -18.73 | 7.41 |
| Right precentral | 13216 | 15.46 | -19.61 | 6.54 | 19661 | 15.03 | -18.77 | 7.06 |
| Left SMA | 18230 | -12.63 | -15.38 | 9.60 | 28033 | -12.93 | -15.96 | 8.46 |
| Right SMA | 17926 | 12.07 | -15.03 | 9.58 | 26484 | 13.13 | -15.80 | 8.13 |
| Left postcentral | 10593 | -17.21 | -22.84 | 6.23 | 20574 | -16.22 | -21.72 | 7.22 |
| Right postcentral | 8764 | 17.33 | -22.24 | 6.20 | 16850 | 16.33 | -21.29 | 7.26 |
| Left Dentate | 1127 | 16.17 | -16.16 | 1.13 | 3192 | 9.46 | -11.11 | -1.31 |
| Right Dentate | 744 | -16.67 | -17.06 | 0.52 | 3028 | -9.34 | -12.24 | -1.77 |

### Variability across individualized connectivity maps

The spatial relation between putative Vim MPMs and their individualized counterpart in a randomly selected, sample subject is depicted in Figure 5 while those of the remaining clusters are provided in Supplementary Figure 3.

**Figure 5.**
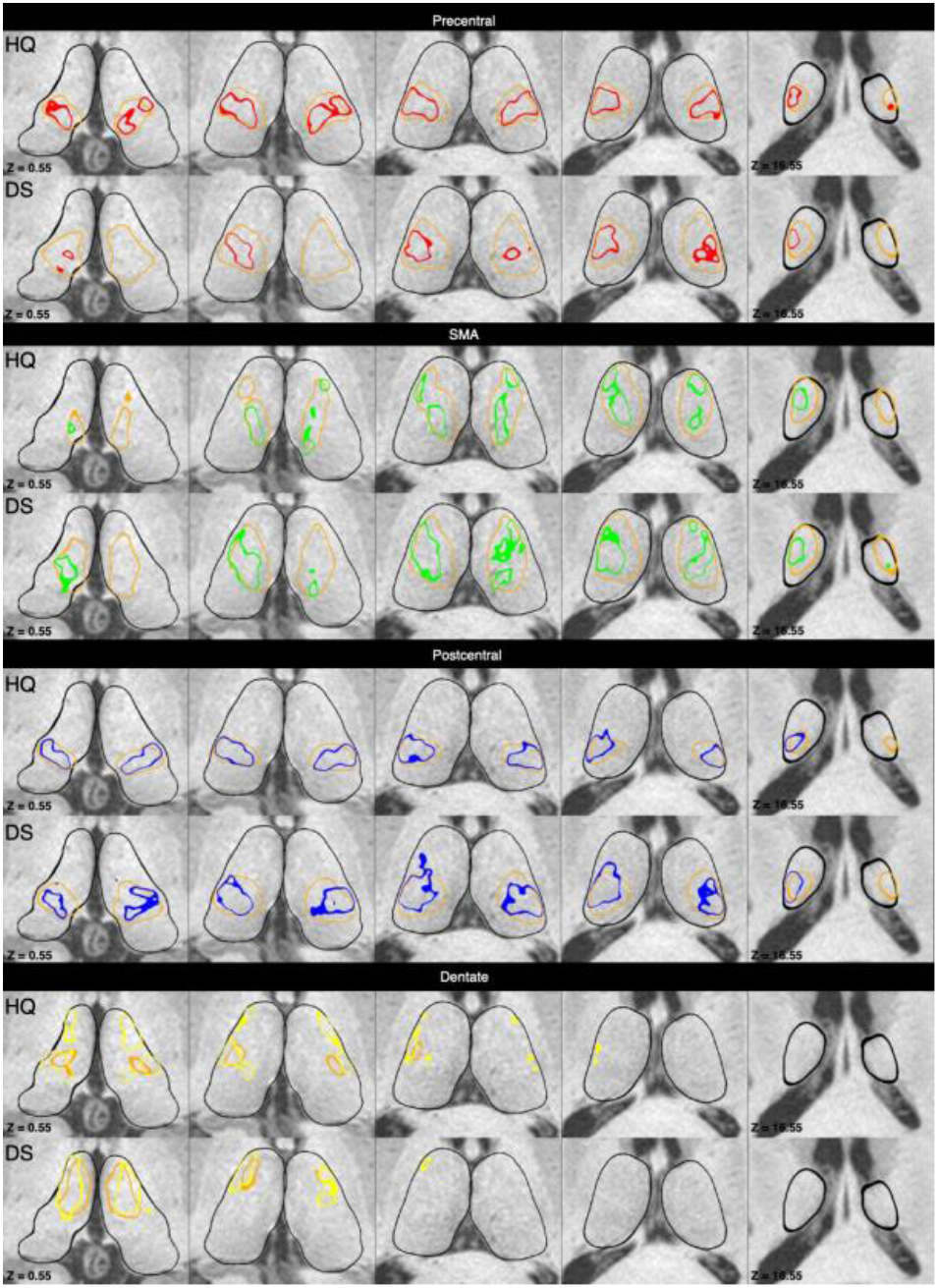
Comparison between tractographic atlas-based and personalized procedures. This figure shows the spatial relations between sensorimotor MPMs registered on a random subject (100408 from the 100 unrelated subjects dataset) and the corresponding individualized connectivity clusters. Multiple axial slices centered on the thalamic region show the MPMs registered on subject native space (light orange) and the corresponding connectivity maps of precentral (red), SMA (green) and dentate (yellow) parcels. For each cluster, the upper rows show the connectivity maps obtained from high quality (HQ) data, while the lower rows display those obtained by the downsampled (DS) dataset.

Average Dice coefficients and Euclidean distances between each selected MPMs and the corresponding individualized maps for both datasets are reported in Table 5. On the other hand, Dice coefficients and Euclidean distances for the remaining MPMs are provided in Supplementary Table 2. Dice coefficients indicated varying similarity between group level and individualized maps for both high quality (range 0.19-0.69) and downsampled (range 0.38-0.70) datasets.

**Table 5.** Average Dice coefficents and Euclidean distances between group-level and individualized sensorimotor maps, in high quality and downsampled datasets. MPMs and individual maps are both referred the most reliable pipeline (CSD-THR).

| Clusters | High Quality |  |  |  | Downsampled |  |  |  |
| --- | --- | --- | --- | --- | --- | --- | --- | --- |
|  | AvgDICE |  | Avg Euclidean Distance |  | Avg DICE |  | Avg Euclidean Distance |  |
|  | Left | Right | Left | Right | Left | Right | Left | Right |
| **Precentral** | 0.65 | 0.69 | 1.49 | 1.31 | 0.66 | 0.66 | 1.24 | 1.44 |
| **SMA** | 0.67 | 0.65 | 1.40 | 1.50 | 0.66 | 0.67 | 1.20 | 1.14 |
| **Postcentral** | 0.65 | 0.63 | 1.65 | 1.67 | 0.63 | 0.63 | 2.26 | 2.14 |
| **Dentate** | 0.23 | 0.19 | 3.30 | 3.54 | 0.41 | 0.38 | 3.67 | 3.77 |

Across high quality datasets, a good spatial correspondence (Dice > 0.5) was observed for connectivity maps, with highest values exhibited by precentral (Left 0.65, Right 0.69) and SMA connectivity clusters (Left 0.67, Right 0.65). Vice versa, the dentate cluster showed the lowest similarity with its group-level counterpart (Left 0.23, Right 0.19). Such trend was maintained across motor-related maps derived from downsampled datasets with higher similarity showed by precentral (Left 0.66, Right 0.66) and SMA (Left 0.66, Right 0.67) parcels; on the other hand, in line with the high-quality dataset, similarity was lower for dentate parcels (Left 0.23, Right 0.19).

Accordingly, the COGs of subject-level clusters derived from connectivity to cortical targets such as precentral (Left 1.49mm, Right 1.31mm) and SMA (Left 1.40mm, Right 1.50mm) exhibited higher spatial proximity to the group-level counterpart, while the COGs of group-level and individualized dentate clusters resulted to be more distant (Left 3.30mm, Right 3.54mm). Similarly, in the downsampled dataset, the COG of atlas-based and individualized precentral (Left 1.24mm, Right 1.44mm) and SMA (Left 1.20mm, Right 1.14mm) parcels exhibited a fair spatial proximity, while those of dentate parcels showed pronounced distance (Left 3.67mm, Right 3.77mm).

Average Euclidean distances between mean COGs of connectivity maps across all individuals and the respective individualized COGs (Figure 6, Table 6) resulted also in remarkable differences ranging from 1.66 to 5.54mm in high quality dataset and from 0.90 to 2.81mm in the downsampled dataset.

**Table 6.** The table summarizes average Euclidean distances between mean COG and individualized COG of the main sensorimotor maps on the ICBM 2009b asymmetric template for both high quality and downsampled datasets.

| Avg Euclidean Distance | High Quality |  | Downsampled |  |
| --- | --- | --- | --- | --- |
|  | Left | Right | Left | Right |
| **Precentral** | 1.69 | 1.79 | 1.10 | 1.17 |
| **SMA** | 1.61 | 1.81 | 1.09 | 1.07 |
| **Postcentral** | 1.80 | 1.89 | 1.27 | 1.31 |
| **Dentate** | 5.23 | 5.54 | 2.73 | 2.81 |

**Figure 6.**
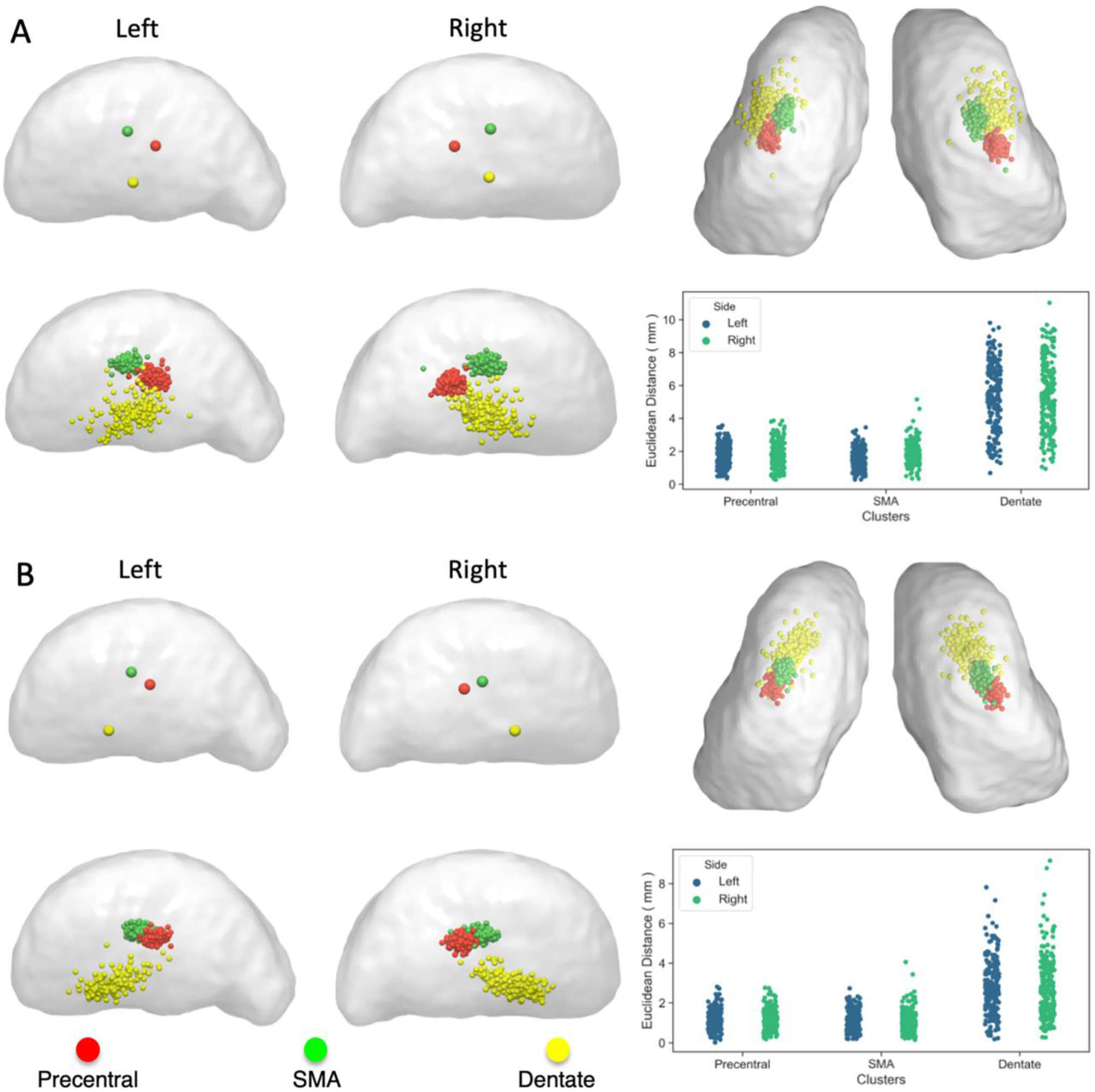
Inter-individual variability. Coordinates of average and individualized COGs of the precentral (red), SMA (green), and dentate (yellow) clusters derived from high quality (A) and downsampled (B) datasets have been mapped on thalamus surfaces. The left half of the figure depicts lateral views of left and right thalamic surfaces on ICBM 2009b template. The right half of the figure shows superior views of thalamic surfaces (top) and strip plots (bottom) reporting Euclidean distances between mean and individual COGs.

The closest spatial relationship between the average COGs and individualized COGs has been observed for precentral (Left 1.69mm, Right 1.79mm) and SMA parcels (Left 1.61mm, Right 1.81) in high quality datasets and for SMA clusters (Left 1.09mm, Right 1.07mm) in downsampled datasets. Conversely, higher Euclidean distances have been observed for dentate clusters both in high quality (Left 5.23mm, Right 5.54mm) and in downsampled (Left 2.73mm, Right 5.54mm) datasets. Such heterogeneity in COG position of the dentate connectivity map is conspicuous if compared to other parcels regardless the dataset quality considered (Figure 6, Table 6).

### Spatial relations between MPMs and optimal stimulation points

Coordinates of optimal stimulation points for ET showed remarkable differences in terms of location (Figure 7A-B).

**Figure 7.**
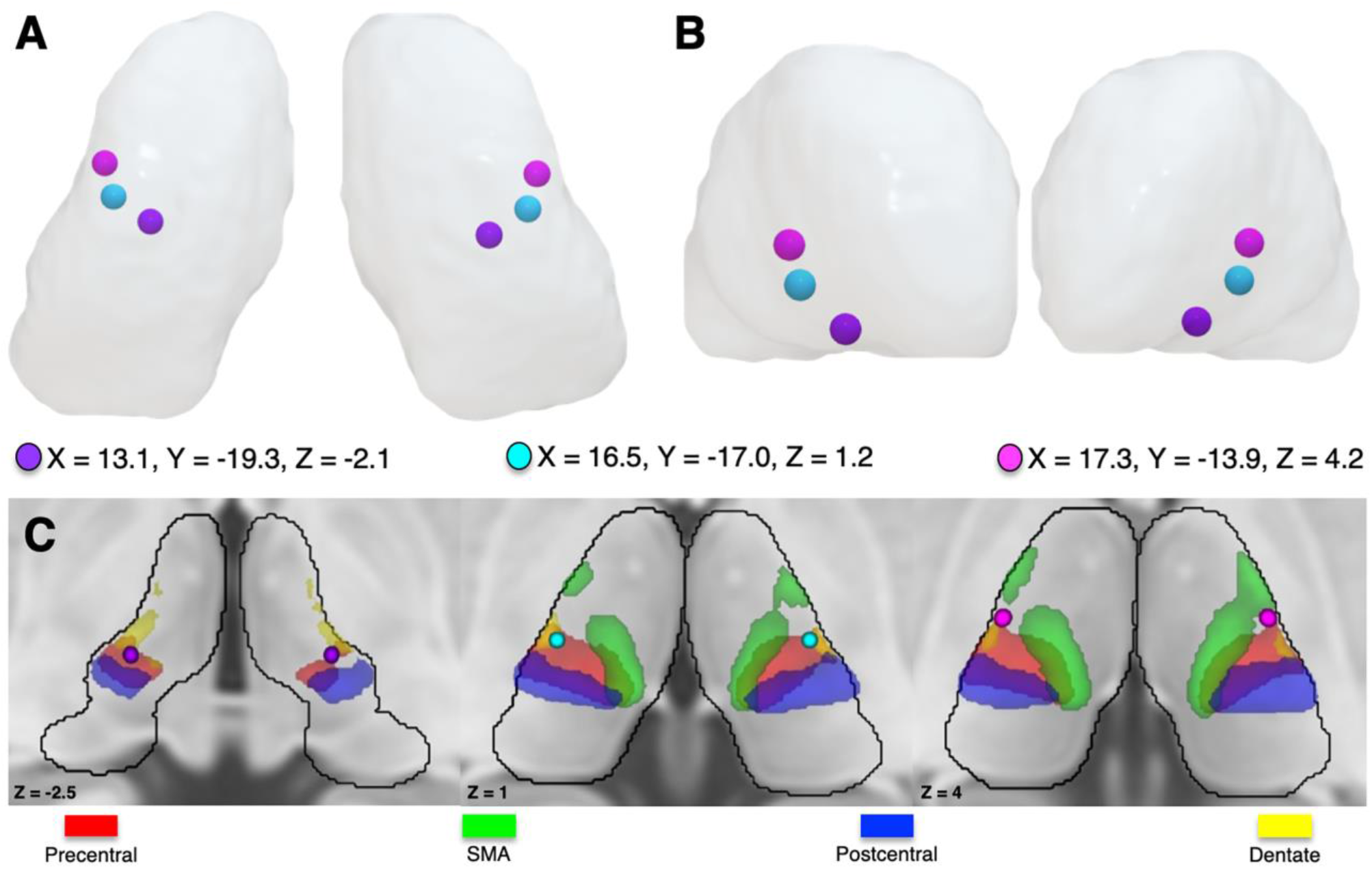
Spatial relations between connectivity maps and optimal stimulation points for essential tremor. The upper part of the figure depicts three stimulation points mapped on thalamus surface in superior (A) and frontal (B) views: the “classic” stimulation point (violet), the voxel most frequently activated in 64 patients implanted with DBS(light blue), the most correlated voxel with clinical improvement (purple). (C) Three axial slices depicting the aforementioned stimulation points and the main sensorimotor MPMs as derived from CSD-THR pipeline.

However, Euclidean distances between COG of MPMs and such literature-based stimulation points (Elias et al., 2020), exhibited the same trend regardless the set of coordinates considered. The classic stimulation point was located in the most ventral thalamic aspect, in close proximity to the posterior subthalamic area and exhibited the highest proximity with the dentate MPMs (Left 5mm, Right 5mm). On the other hand, COG of precentral (Left 7mm, Right 6.4mm), SMA (Left 12.3mm, Right 9.6mm) and postcentral (Left 9.4mm, Right 9.8mm) MPMs, showed remarkable Euclidean distances with such coordinates. Stimulation points derived from unweighted frequency maps were located in a more dorsal position and showed overall the lowest Euclidean distance with dentate group-level connectivity maps (Left 0.7mm, Right 0.9mm), whilst higher Euclidean distances were exhibited by COG of precentral (Left 6.7mm, Right 6mm), SMA (Left 9.4mm, Right 9.7mm) and postcentral (Left 9.2mm, Right 8.5mm) MPMs. Finally, coordinates derived clinically weighted hotspots were located even more dorsally and anteriorly, and showed the lowest Euclidean distance with COG of dentate (Left 4mm, Right4.9mm) clusters; conversely, higher distances were found for precentral (Left 7mm, Right 6.4mm), SMA (Left 7.3mm, Right 7.6mm) and postcentral MPM COGs (Left 9.2mm, Right 9.7mm) (Figure 7C, Table 7).

**Table 7.** Table summarizing Euclidean distances (in mm) between COG of MPMs and coordinates of optimal stimulation points for ET.

| Cluster | Optimal stimulation points |  |  |  |  |  |
| --- | --- | --- | --- | --- | --- | --- |
|  | Classical |  | Unweighted |  | Weighted |  |
|  | Left | Right | Left | Right | Left | Right |
| **Precentral** | 7.0 | 6.4 | 6.7 | 6.0 | 4.0 | 4.9 |
| **SMA** | 12.3 | 9.6 | 9.4 | 9.7 | 7.0 | 6.4 |
| **Postcentral** | 9.2 | 8.5 | 9.2 | 8.5 | 7.3 | 7.6 |
| **Dentate** | 5.0 | 5.0 | 0.7 | 0.9 | 9.2 | 9.7 |

### Effects of pipelines and data quality on connectivity clusters

For each putative Vim cluster (precentral, SMA, dentate), F-values describing main effects and interactions of data quality and parcellation pipelines on SDI, are reported below. A Mauchly’s test was carried out to verify sphericity of data. When sphericity assumption was not met degrees of freedom and their related error was Greenhouse-Geisser corrected. Boxplots summarizing the distribution of SDI for each connectivity cluster are provided in Figure 8. Results concerning the remaining parcels (paracentral, postcentral, prefrontal, parietal, temporal, occipital) are included in Supplementary Tables 4-6.

**Figure 8.**
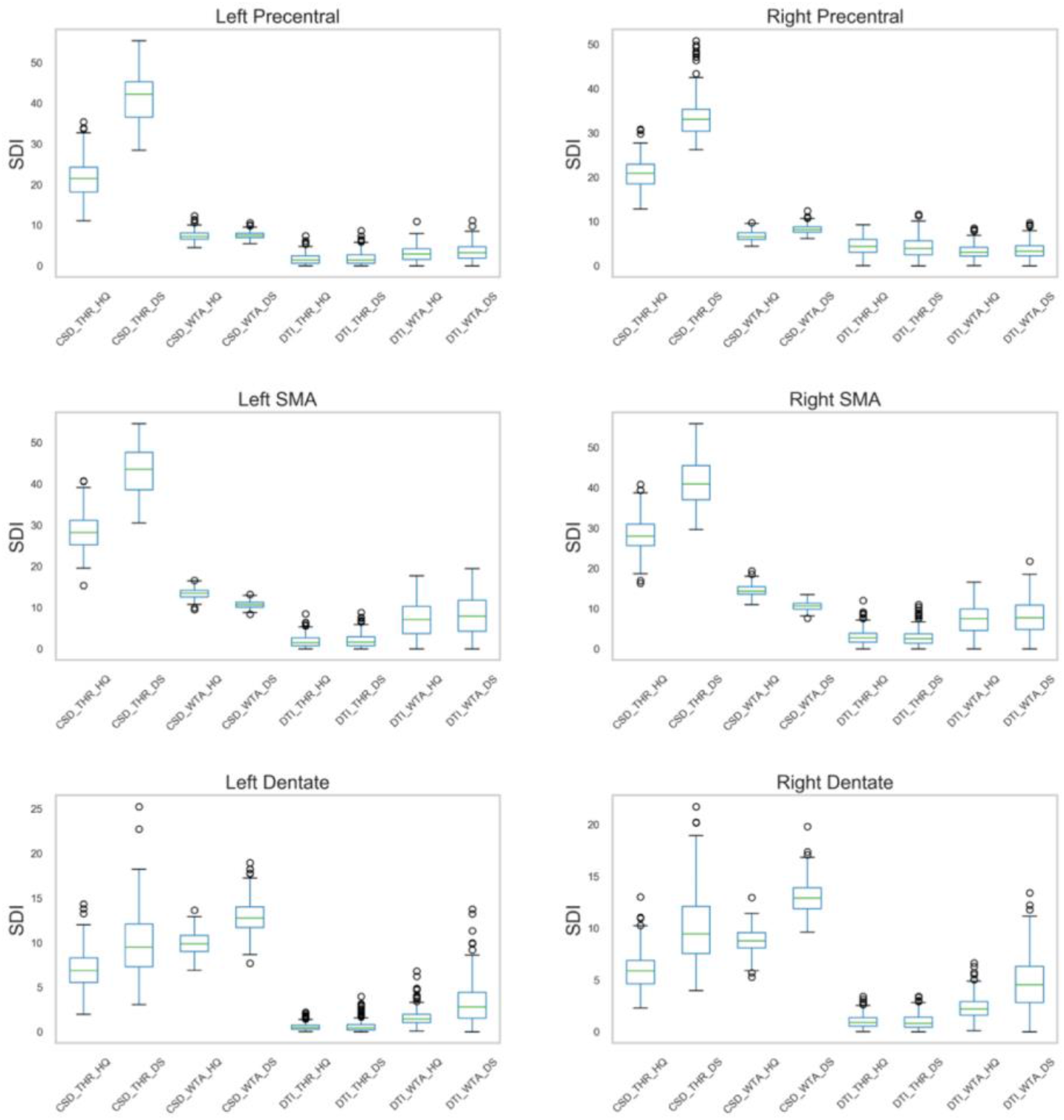
Connectivity clusters quantitative analysis. Boxplots showing the distribution of streamline density index (SDI), grouped for datasets and pipelines, for sensorimotor clusters which have been related to tremor treatment in existing literature. Boxes refer to quartiles of SDI values, their extension is equal to the standard deviation from the median SDI value (horizontal line); whiskers depict SDI values which fall out of from standard deviation. Outliers are shown as blank circles.

For left precentral clusters, we found a significant main effect of data quality (F(1,209)= 2837.815, p < .001, 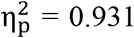) and pipelines (F(1,17, 275.301)= 6892.535, p < .001, 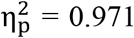) on SDI. In particular, downsampled datasets displayed significantly higher SDI if compared to high quality ones (p < .001) whilst different pipelines showed significantly different pipelines with CSD-based pipelines showing the highest volumes (p < .001, Bonferroni corrected). We observed also a significant interaction between data quality and pipeline (F(1.208, 252.421)= 2791.503, p < .001, 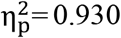). A similar pattern was observed for right precentral clusters, which exhibited significant main effect for both data quality (F(1,209)=1667.440, p < .001, 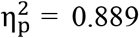) and pipelines (F(1.826, 381.537)= 6921.534, p < .001, 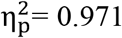) with higher SDI across downsampled datasets (p < .001) and significant differences across parcellation pipelines (p < .001, Bonferroni corrected). A significant interaction was noticed between data quality and pipelines (F(1.268,265.069)=1284.393, p < .001, 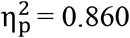).

Also left SMA connectivity maps exhibited a significant main effect of data quality (F(1,209)= 1025.421, p < .001, 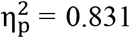) and pipelines (F(1.764)= 5073.377, p < .001, 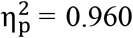) on SDI values, with downsampled datasets exhibited higher SDI than high quality ones (p < .001) and higher SDI across CSD-based pipelines (p < .001, Bonferroni corrected). Moreover, we noticed a significant interaction between data quality and pipelines (F(1.475, 308.321)= 1303.401, p < .001, 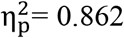) such that data quality influenced mostly CSD-THR pipeline (F(1,209)= 1599.236, p< .001, 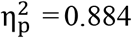) and CSD-WTA (F(1,209)= 925.134, p< .001, 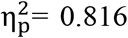), pipelines. Similarly right SMA connectivity maps showed significant main effect for data quality (F(1,209)= 506.150, p < .001, 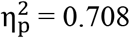) and pipelines (F(2.149,449.039)= 5620.387, p < .001, 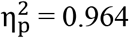) with downsampled datasets showing significantly higher SDI than high quality ones (p < .001) and CSD-based pipelines resulting in higher SDI values than DTI-based ones (p< .001, Bonferroni corrected). Interaction between data quality and pipelines was significant (F(1.456, 304.323)= 1086.817, p < .001, 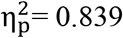).

A significant main effect of both data quality (F(1,209)= 364.643, p < .001, 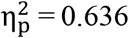) and pipelines (F(1.724,360.293)= 3221.193, p < .001, 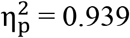) was observed also for left dentate clusters, with downsampled datasets showing higher SDI than high quality ones (p< .001) and CSD-WTA pipelines resulting in clusters characterized by highest SDI values (p < .001, Bonferroni corrected). Moreover we noticed a significant interaction between data-quality and pipelines (F(1.555, 325.008)= 97.647, p < .001, 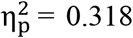). The same trend was exhibited for right dentate clusters: both data quality (F(1,209)= 886.333, p < .001, 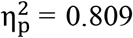) and pipelines (F(1.763, 368.505)= 2471.665, p < .001, 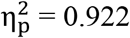) showed a significant main effect with downsampled datasets showing higher SDI values (p < .001) and pipelines displaying different effect on SDI, with the highest values reported for CSD-based pipelines (p < .001, Bonferroni corrected). The interaction between data quality and pipeline was significant (F(1.802, 376.565)= 237.181, p < .001, 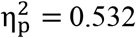).

## Discussion

In the last decades, connectivity-derived thalamic parcellation has been accomplished using different imaging modalities both with hypothesis-driven and data-driven methods (Behrens et al., 2003; Fan et al., 2015; Lambert et al., 2017; O’muircheartaigh et al., 2015; Zhang et al., 2008). While hypothesis driven approaches require arbitrary selection of the targets connected to the structure to be parcellated (Cacciola et al., 2019a; Patriat et al., 2018; Plantinga et al., 2018), data-driven approaches rely on different clustering algorithms to subdivide anatomical structures in a pre-defined number of clusters (Eickhoff et al., 2015; Saygin et al., 2011).

In the present study we tested a traditional hypothesis-driven CBP of thalamus on a large sample of healthy subjects from the HCP. Despite the known limitations of the hypothesis driven approach, including the introduction of a selection bias regarding the pre-selected target regions, our choice was motivated by the need of decreasing the number of variables which may affect the results of CBP. Indeed, data-driven methods strongly rely on the selected clustering algorithm, the chosen number of clusters and the validation metrics for evaluating stability of clustering solution (Reuter et al., 2020). Moreover, establishing anatomical priors may constitute a strength in stereotactic neurosurgery where anatomy and pathophysiology are strongly tied together in disease-specific models which drive the choice of targets (Helmich et al., 2013; Middlebrooks et al., 2020). Finally, hypothesis-driven methods are more straightforward and easier to implement in clinical practice for preoperative evaluation or retrospective analysis as already demonstrated by existing studies (Patriat et al., 2018; Plantinga et al., 2018).

Given the heterogeneity in acquisition parameters, diffusion modelling techniques and voxel classification criteria in existing studies, we combined the most commonly employed diffusion signal modelling techniques (CSD and DTI) and voxel classification criteria (THR-based and WTA approaches) thus resulting into four parcellation pipelines (i.CSD-THR ii.CSD-WTA iii. DTI-THR iv. DTI-WTA) for each subject. With the aim of identifying a parcellation pipeline that could work either on state-of-art data as well as on standard quality, clinical-like MRI acquisitions, we downsampled high-quality structural data from the HCP repository in order to match the most relevant features of DWI (voxel size and directions) that are commonly acquired in advanced-care clinical contexts. The choice of downsampling previously existing data, instead of directly testing our pipeline on lower-level acquisitions, would ensure that the high- and low-quality datasets would share exactly the same amount of variance (e.g. removing the effects of inter-subject anatomical variability, subject motion or susceptibility-induced artifacts) and that differences in between-subjects similarity could be more easily attributed to the different DWI data quality.

As a first selection step, we conducted an inter-subject similarity analysis to identify a pipeline which leads to more reliable results across subjects. Our results exhibited fair inter-subject reliability both at parcellation (TAO) and clusters level (OBL), showing similar values, even if slightly lower than those reported by a previous analysis (Traynor et al., 2010). This finding could be explained by the larger sample considered in this work (n=210 vs n=18) which likely implies more inter-individual variability. The higher values obtained by CSD-THR pipeline both in high quality and downsampled datasets, strongly suggest that, even in presence of low spatial and angular resolution, connectivity maps obtained using such approach could maintain a similar level of spatial coherence across subjects. By contrast, CSD-WTA pipeline showed fair reliability when applied in high quality datasets, but notably lower values when the same pipeline was implemented on downsampled datasets, thus suggesting more sensitivity to data quality; DTI-based pipelines showed low reliability with minimal differences between high quality and downsampled datasets.

Voxel classification criteria to be employed in the context of hypothesis-driven parcellation remain an open matter of debate; hard-segmentation has been criticized for the assumption of attributing univocally a parcel to a predefined connectivity pattern when it may actually contain multiple fiber populations (Patriat et al., 2018; Sudhyadhom et al., 2013). Caution is needed when interpreting results derived from hard-segmentation, as it has been demonstrated that apparently meaningful parcellations may be retrieved also from extremely corrupted data (Clayden et al., 2019). While the authors suggest that the use of threshold-based approaches could overcome this problem, by filtering out voxels characterized by minor tract density that are more likely to result from spurious connectivity, on the other hand, such procedure requires the choice of arbitrary thresholds which may actually lead to underestimation or overestimation of the results (Middlebrooks et al., 2018c; Plantinga et al., 2018). Our results suggest that, when dealing with higher-order signal modeling such as CSD, the choice of winner-takes-all or threshold-based approach as a final step of the parcellation protocol may have a relevant impact on between-subjects similarity measures, which is likely to be exacerbated when higher-order signal modeling is applied to low spatial and angular resolution datasets. Indeed, when CSD signal modelling is applied on clinical-like datasets, the resulting connectivity parcels may result to be highly variable across subjects because of the higher number of false-positive and spurious streamlines (Essayed et al., 2017). Conversely, DTI-based pipelines show substantially lower reliability regardless of data quality and seems to benefit from WTA in terms of between-subject similarity of spatial maps.

### Spatial relations between connectivity-derived sensorimotor maps, histological Vim and optimal stimulation points

Once the most reliable approach was identified both for high-quality and downsampled datasets, we computed the Dice coefficient between individualized connectivity maps and a histology-derived map of Vim to investigate which connectivity pattern would best suit as a proxy for connectivity-derived Vim identification. Several neurophysiological studies underlined that abnormal oscillations across the DRTC tract may play an important role in the pathophysiology of tremors (Deuschl et al., 1996; Schnitzler et al., 2009), and that beneficial effects of DBS may be linked to the stimulation of DRTC tract or the caudal zona incerta (Akram et al., 2018; Al-Fatly et al., 2019; Blomstedt et al., 2018), suggesting that regions where DRTC tract intersects the thalamus may putatively correspond to the Vim (Fiechter et al., 2017). In addition, since the Vim is commonly defined as a region mainly receiving cerebellar afferent connections and projecting to primary motor cortex (Asanuma et al., 1983a, 1983b), neuroimaging studies proposed thalamic regions connected to precentral gyrus, or to its somatotopic subdivisions (the hand-knob region), as potentially corresponding to such histological entity (Tian et al., 2018; Tsolaki et al., 2018). Finally, other studies proposed also thalamic regions connected to SMA/premotor cortices as functional targets possibly corresponding to ventro-oralis posterior (Vop) nucleus which receives projections from the GPi (Middlebrooks et al., 2018b; Morel et al., 1997; Pouratian et al., 2011).

Our results showed that, in line with the anatomical definition, the highest similarity with the histological Vim was showed by the dentate and precentral connectivity clusters, which ideally correspond to afferent and efferent projection areas of the Vim, respectively (Morel et al., 1997). On the other hand, very poor similarity was observed with postcentral, parietal, temporal and occipital connectivity clusters in both high-quality and clinical-like datasets.

However, it should be noted that low Dice values (< 0.5) were demonstrated by all motor-related connectivity clusters. The low similarity may be motivated by the fact that connectivity-based maps differ from cytoarchitectonic nuclei in size, shape and orientation (Ilinsky et al., 2018), therefore, tractography-derived functional targets are likely to straddle cytoarchitectonic boundaries leading to the observed morphological differences (Akram et al., 2018). Indeed, it should be kept in mind that the nomenclature of thalamic subdivisions is mainly based on histochemical staining of serial sections, rather than structural/functional connectivity (Morel et al., 1997). In addition, the histology-based Vim chosen for this analysis is derived from a single post-mortem specimen, which is unlikely to perfectly match the anatomy of the wider sample (n=210) here considered.

Beyond comparing the spatial distribution between connectivity-derived maps and anatomy of Vim thalamic nucleus defined as an histological entity, we also investigated the spatial relations of the group-level connectivity maps with higher similarity to Vim with three different, group-level coordinates of optimal stimulation points for ET, which have been recently provided in a study conducted on implanted patients (Elias et al., 2020) (Figure 8). Specifically, the first stimulation site consisted of the “classic” electrode location, commonly used to guide stereotactic neurosurgery of tremor (Horn et al., 2017; Papavassiliou et al., 2004), whilst the second and the third sets of coordinates have been obtained from probabilistic stimulation mapping (PSM) of volume of tissue activated (VTA), representing the most commonly stimulated voxels in the cohort (clinically unweighted maps) and clinically-weighted hotspots (zones where stimulation produced above-mean clinical improvements) respectively (Elias et al., 2020). “Classic” coordinates of electrode location were located in the ventral-most part of thalamus, just adjacent to dentate and precentral maps; such position likely represents the best match to the classic localization of Vim, as described in anatomical studies (Asanuma et al., 1983c, 1983a). The most stimulated voxel in the cohort (clinically unweighted maps) was located more dorsally, being entirely centered in the dentate MPMs. Finally, the clinically-weighted hotspot was located further dorsally and anteriorly, lying just anterior to dentate and precentral MPMs encroaching SMA connectivity maps, as suggested by other existing studies (Middlebrooks et al., 2018b; Pouratian et al., 2011).

Interestingly, the spatial relations here described agree with data-driven connectomic analyses provided by normative connectomes on ET implanted patients (Al-Fatly et al., 2019; Elias et al., 2020). In the study conducted by Al-Fatly and colleagues structural and functional “therapeutic” connectivity patterns of leads located in close proximity to “classic” coordinates included cerebellum, pre and postcentral gyri, SMA and paracentral lobule; moreover, discriminative fiber tracts connected to VTA, thus being linked with clinical improvement, belong to DRTC tract (Al-Fatly et al., 2019; Baldermann et al., 2019). Similarly, in the study conducted by Elias and collaborators, clinically-weighted hotspots were connected to precentral gyrus, SMA and prefrontal regions, including again the DRTC (Elias et al., 2020). Herein, we show that the classic stimulation point lies in close proximity to dentate, precentral, SMA and paracentral MPMs. It may be hypothesized that DBS of DRTC entry zone, may exert a modulatory influence on neighbouring thalamic structures characterized by connectivity with those areas. Accordingly, we observed spatial proximity of the clinically weighted hotspots with precentral and SMA, but a slight anterior displacement from dentate MPMs.

Euclidean distances between COGs of the sensorimotor MPMs and coordinates of stimulation points suggest that dentate connectivity maps exhibit the most pronounced spatial proximity with each of the coordinates considered. Even if such finding is consistent with studies suggesting the DRTC as the main target for tremor-treatment and on discriminative fibers tracts analysis conducted on implanted patients, it is important to keep in mind that COG largely depends on the shape of connectivity clusters and does not necessarily correspond to the optimal location for stimulation delivery within connectivity maps, as it needs to be addressed by further studies on patient cohorts. Once we selected the three spatial connectivity patterns that are more reliable in terms of inter-subject variability for connectivity-based Vim identification, we sought to characterize both similarity and COG spatial proximity between subject-level and group-level maps, attempting to get a measure of how much the group-level maps obtained in the present study are representative of inter-individual differences. Hence, we registered tractography-derived MPMs to each subject native space and computed the Dice coefficients between each pair of images and the Euclidean distance between the COG of MPMs and their individualized counterpart.

We found overall high similarity for all connectivity parcels related to cortical targets in both high quality and downsampled datasets, while, conversely, dentate connectivity maps exhibited the lowest Dice coefficient and highest between-COG Euclidean distances. These results are in line with those provided by Akram and colleagues which studied the spatial relations between group-level dento-thalamic connectivity maps and patient based maps, finding large Euclidean distances between group-level and individualized maps (Akram et al., 2018). Summarizing, spatial maps derived from dento-thalamic connectivity seems to be more influenced by inter-individual variability, suggesting that group-level maps could be, in this case, less representative of individual connectivity patterns.

We further analyzed inter-individual variability of these connectivity parcels, by assessing Euclidean distances between each subject’s COG and the COG averaged across all subjects. Whilst spatial proximity was observed for almost all connectivity clusters, including those that may have implications in functional neurosurgery of tremor such as precentral and SMA maps (Middlebrooks et al., 2018b; Tian et al., 2018; Tsolaki et al., 2018), we found the highest Euclidean distances for the dentate clusters. Such variability may be due to technical reasons, as the DRTC is a long, decussating tract, often hard to reconstruct using tractography (Sinke et al., 2018), despite the reconstruction strategy applied herein has been already tested in other studies (Palesi et al., 2016, 2015; Tang et al., 2018). Another possible hypothesis is that the observed displacement may represent actual inter-individual anatomical variability, as demonstrated both in ex-vivo in thalamic nuclear boundaries (Morel et al., 1997) and in-vivo in putative Vim identified using tractography and functional MRI (Anderson et al., 2011; Anthofer et al., 2014; Kincses et al., 2012). Such inter-individual variability may also explain the remarkable heterogeneity of clinical outcomes, with improvement rates ranging from 18 to 88%, observed in patients who underwent Vim DBS via traditional atlas-based targeting (Middlebrooks et al., 2018a).

### Data quality and pipelines impacts the volumes of connectivity-derived maps

Finally, we aimed at quantifying the impact of methodological variables on the volumetric estimates of the resulting connectivity maps, by evaluating the effects of data quality and pipelines over a volume-based metric, namely the SDI (Theisen et al., 2017).

We found significant differences between clusters obtained using CSD and DTI diffusion signal modeling with the former resulting in significantly higher SDI values. This result may be explained by the intrinsic properties of CSD signal modeling which resolves complex fibers configuration leading to more exhaustive reconstruction of white matter bundles (Barbeau et al., 2020). On the other hand, it is likely that modelling only principal diffusion direction entails an underestimation of the reconstructed tractograms, leading to small and circumscribed connectivity clusters (Farquharson 2013). In addition, we noticed that the choice of voxel classification criteria had a different effect when applied after CSD or DTI-signal modelling: in the former case, higher SDI values were exhibited when an adaptive threshold was applied, in the latter, hard-segmentation led to higher SDI values.

Moreover, significant volumetric differences were noticed between clusters obtained using high quality and downsampled datasets. All clusters derived from downsampled datasets were significantly larger than those derived from high quality ones. Such result could be explained by the poorer spatial and angular resolution in the downsampled sets which lead to an increased noise during tracking process and subsequently to larger and coarser maps overlapping each other (Barbeau et al., 2020). The interaction between data quality and parcellation pipelines was significant across different connectivity parcels suggesting that different combination of data quality and pipelines may contribute differently to variations observed in SDI. Effect sizes suggest that data quality has a higher influence on SDI values pertaining to CSD-based pipelines rather than DTI-based ones. While being present, such difference between CSD-and DTI-based parcellation pipelines due to data quality is mitigated for the dentate clusters suggesting greater sensibility to data quality when complex-configuration bundles are reconstructed using DTI signal modelling. On the other hand, volumetric differences related to the use of different pipelines resulted to be always remarkable regardless data quality. Such result may suggest a prominent role of pipeline choice over data quality in CBP settings.

## Limitations

This study has several limitations that need to be addressed. Firstly, tractography cannot distinguish polarity of white matter bundles, detect synapses or reconstruct intracortical white matter (Jbabdi and Johansen-Berg, 2011). However, the main purpose of this study was to test impact of different methodologies on an anatomically informed parcellation scheme (Behrens et al., 2003; Johansen-Berg et al., 2005; Traynor et al., 2010) rather than characterize exhaustively thalamo-cortical and cortico-thalamic connections, their origin from thalamic histologic subdivisions or their termination within cortical sheet.

We acknowledge that connectivity maps obtained from CBP may be more properly defined as “connectivity modules” within parcellated structures but do not biologically imply that those effective parcellation exists within the structures considered. As a matter of fact, it has been underlined that connectivity-based thalamic parcellation probably underestimates the nuclear thalamic complexity (Iglesias et al., 2018; Su et al., 2019). Even if the overall organization here described agrees with the economy of brain wiring (Van Essen, 1997) and tractography-derived maps have been somehow reconducted to groups of thalamic nuclei (Behrens et al., 2003), this study is not aimed at distinguishing thalamic cytoarchitectonic subdivisions. Indeed, connectivity methods may identify a functional target which is only putatively reconducted to the Vim or to other thalamic nuclei to maintain adherence with available pathophysiological models (Helmich et al., 2013). Nonetheless, in-vivo reconstruction of white matter has been demonstrated to be efficient in recognizing functional targets in an individualized fashion, improving clinical outcomes (Fenoy and Schiess, 2018).

Our results pointed out that a more reliable parcellation may be obtained using threshold-based approach, in agreement with previous studies showing, by means of preliminary electrophysiology data, that position of threshold-based connectivity clusters agrees well with underlying neuronal populations activity of different thalamic nuclei (Elias et al., 2012; Sudhyadhom et al., 2013).

It is worth to note that we evaluated reliability by assessing similarity between connectivity parcels across subjects belonging to the same sample, and cannot therefore disentangle the effect of anatomical variability of structures of interest from reliability of the parcellation pipeline. However, since we employed the same sample for the entire analysis, we may assume inter-subject anatomical variability as constant and interpret differences in similarity as a consequence of parcellation pipelines.

Finally, we cannot recommend any of the proposed connectivity map as a better stimulation target in respect of the others, since such assumption need to be corroborated by clinical data on patients’ cohorts. However, we believe that MPMs based on a large cohort may further inform the interpretation of outcomes of both retrospective and prospective studies, especially if joined with data-driven connectomic analysis methods.

## Conclusions

In this work we investigated the influence of main methodological variables when performing individualized thalamic-CBP for possible applications in functional neurosurgery settings. We found that applying a pipeline that involves CSD-signal modelling and a threshold-based approach provides fair reliable estimates in terms of inter-subject variability of most motor-related thalamic parcels both in high quality and in downsampled, clinical-like, MRI acquisition. However, it is important to bear in mind that thalamic dMRI connectivity-based segmentation should be carefully checked in stereotactic targeting where a good outcome hinges on millimetric accuracy. Indeed, our findings further confirm that structural connectivity-derived maps differ for neuroanatomical orientation, shapes, and sizes when compared to a ground truth model based on histochemical staining on seriate sections of human brain. In addition, we found that DRTC tract, as reconstructed by diffusion tractography, is highly variable across subjects and its anatomical plausibility need to be carefully verified if used prospectively to guide neurosurgical targeting. Finally, we observed noteworthy effects of data quality and parcellation pipelines on connectivity clusters volumes, highlighting major inconsistencies that can derived by the high variability in dMRI acquisition and processing.

We hope that this work, will be of interest for further research focusing on advanced MRI based techniques for tailoring treatment on individuals and interpret outcomes using connectomic analyses.

## Acknowledgements

Data were provided by the Human Connectome Project, WU-Minn Consortium (Principal Investigators: David Van Essen and Kamil Ugurbil; 1U54MH091657), funded by the 16 NIH institutes and centers that support the NIH Blueprint for Neuroscience Research; and by the McDonnell Center for Systems Neuroscience at Washington University.

## Author statement

GAB: Conceptualization, Investigation, Writing – original draft; Writing – review & editing, Visualization; SB: Conceptualization, Investigation, Writing – original draft; Writing – review & editing, Visualization; AB: Resources, Data curation; RC: Resources, Data curation; AT: Resources, Data curation, Writing – original draft; GA: Resources, Data curation, Writing – review & editing; DM: Conceptualization, Investigation, Supervision, Writing – original draft; Writing – review & editing; AC: Conceptualization, Investigation, Project administration, Supervision, Writing – original draft; Writing – review & editing.

## Data availability

Data were provided by the Human Connectome Project, WU-Minn Consortium (Principal Investigators: David Van Essen and Kamil Ugurbil; 1U54MH091657) and are publicly available at https://www.humanconnectome.org/study/hcp-young-adult/document/1200-subjects-data-release.

## References

Akram, H., Dayal, V., Mahlknecht, P., Georgiev, D., Hyam, J., Foltynie, T., Limousin, P., De Vita, E., Jahanshahi, M., Ashburner, J., Behrens, T., Hariz, M., Zrinzo, L., 2018. Connectivity derived thalamic segmentation in deep brain stimulation for tremor. NeuroImage Clin. https://doi.org/10.1016/j.nicl.2018.01.008

Akram, H., Hariz, M., Zrinzo, L., 2019. Connectivity derived thalamic segmentation: Separating myth from reality. NeuroImage Clin. https://doi.org/10.1016/j.nicl.2019.101758

Al-Fatly, B., Ewert, S., Kübler, D., Kroneberg, D., Horn, A., Kühn, A.A., 2019. Connectivity profile of thalamic deep brain stimulation to effectively treat essential tremor. Brain. https://doi.org/10.1093/brain/awz236

Ambrosen, K.S., Eskildsen, S.F., Hinne, M., Krug, K., Lundell, H., Schmidt, M.N., van Gerven, M.A.J., Mørup, M., Dyrby, T.B., 2020. Validation of structural brain connectivity networks: The impact of scanning parameters. Neuroimage 204. https://doi.org/10.1016/j.neuroimage.2019.116207

Anderson, J.S., Dhatt, H.S., Ferguson, M.A., Lopez-Larson, M., Schrock, L.E., House, P.A., Yurgelun-Todd, D., 2011. Functional connectivity targeting for deep brain stimulation in essential tremor. Am. J. Neuroradiol. 32, 1963–1968. https://doi.org/10.3174/ajnr.A2638

Anthofer, J., Steib, K., Fellner, C., Lange, M., Brawanski, A., Schlaier, J., 2014. The variability of atlas-based targets in relation to surrounding major fibre tracts in thalamic deep brain stimulation. Acta Neurochir. (Wien). https://doi.org/10.1007/s00701-014-2103-z

Asanuma, C., Thach, W.T., Jones, E.G., 1983a. Distribution of cerebellar terminations and their relation to other afferent terminations in the ventral lateral thalamic region of the monkey. Brain Res. Rev. https://doi.org/10.1016/0165-0173(83)90015-2

Asanuma, C., Thach, W.T., Jones, E.G., 1983b. Anatomical evidence for segregated focal groupings of efferent cells and their terminal ramifications in the cerebellothalamic pathway of the monkey. Brain Res. Rev. https://doi.org/10.1016/0165-0173(83)90016-4

Asanuma, C., Thach, W.T., Jones, E.G., 1983c. Cytoarchitectonic delineation of the ventral lateral thalamic region in the monkey. Brain Res. Rev. https://doi.org/10.1016/0165-0173(83)90014-0

Avants, B., Tustison, N., Song, G., 2009. Advanced Normalization Tools (ANTS). Insight J.

Baldermann, J.C., Melzer, C., Zapf, A., Kohl, S., Timmermann, L., Tittgemeyer, M., Huys, D., Visser-Vandewalle, V., Kühn, A.A., Horn, A., Kuhn, J., 2019. Connectivity Profile Predictive of Effective Deep Brain Stimulation in Obsessive-Compulsive Disorder. Biol. Psychiatry. https://doi.org/10.1016/j.biopsych.2018.12.019

Barbeau, E.B., Descoteaux, M., Petrides, M., 2020. Dissociating the white matter tracts connecting the temporo-parietal cortical region with frontal cortex using diffusion tractography. Sci. Rep. 10, 1–13. https://doi.org/10.1038/s41598-020-64124-y

Basser, P.J., Mattiello, J., Lebihan, D., 1994. Estimation of the Effective Self-Diffusion Tensor from the NMR Spin Echo. J. Magn. Reson. Ser. B. https://doi.org/10.1006/jmrb.1994.1037

Behrens, T.E.J., Berg, H.J., Jbabdi, S., Rushworth, M.F.S., Woolrich, M.W., 2007. Probabilistic diffusion tractography with multiple fibre orientations: What can we gain? Neuroimage 34, 144–155. https://doi.org/10.1016/j.neuroimage.2006.09.018

Behrens, T.E.J., Woolrich, M.W., Smith, S.M., Boulby, P.A., Barker, G.J., Sillery, E.L., Sheehan, K., Ciccarelli, O., Thompson, A.J., Brady, J.M., Matthews, P.M., 2003. Non-invasive mapping of connecitons between human thalamus and cortex using DTI. Nat. Neurosci.

Benabid, A.L., Pollak, P., Gao, D., Hoffmann, D., Limousin, P., Gay, E., Payen, I., Benazzouz, A., 1996. Chronic electrical stimulation of the ventralis intermedius nucleus of the thalamus as a treatment of movement disorders. J. Neurosurg. https://doi.org/10.3171/jns.1996.84.2.0203

Bertino, S., Basile, G.A., Bramanti, A., Anastasi, G.P., Quartarone, A., Milardi, D., Cacciola, A., 2020. Spatially coherent and topographically organized pathways of the human globus pallidus. Hum. Brain Mapp. 41, 4641–4661. https://doi.org/10.1002/hbm.25147

Blomstedt, P., Persson, R.S., Hariz, G.M., Linder, J., Fredricks, A., Häggström, B., Philipsson, J., Forsgren, L., Hariz, M., 2018. Deep brain stimulation in the caudal zona incerta versus best medical treatment in patients with Parkinson’s disease: A randomised blinded evaluation. J. Neurol. Neurosurg. Psychiatry 89, 710–716. https://doi.org/10.1136/jnnp-2017-317219

Boutet, A., Gramer, R., Steele, C.J., Elias, G.J.B., Germann, J., Maciel, R., Kucharczyk, W., Zrinzo, L., Lozano, A.M., Fasano, A., 2019. Neuroimaging Technological Advancements for Targeting in Functional Neurosurgery. Curr. Neurol. Neurosci. Rep. https://doi.org/10.1007/s11910-019-0961-8

Broser, P., Vargha-Khadem, F., Clark, C.A., 2011. Robust subdivision of the thalamus in children based on probability distribution functions calculated from probabilistic tractography. Neuroimage 57, 403–415. https://doi.org/10.1016/j.neuroimage.2011.04.054

Cacciola, A., Milardi, D., Basile, G.A., Bertino, S., Calamuneri, A., Chillemi, G., Paladina, G., Impellizzeri, F., Trimarchi, F., Anastasi, G., Bramanti, A., Rizzo, G., 2019a. The cortico-rubral and cerebello-rubral pathways are topographically organized within the human rwed nucleus. Sci. Rep. 9, 1–12. https://doi.org/10.1038/s41598-019-48164-7

Cacciola, A., Milardi, D., Bertino, S., Basile, G.A., Calamuneri, A., Chillemi, G., Rizzo, G., Anastasi, G., Quartarone, A., 2019b. Structural connectivity-based topography of the human globus pallidus: Implications for therapeutic targeting in movement disorders. Mov. Disord. 34, 987–996. https://doi.org/10.1002/mds.27712

Calamante, F., Tournier, J.D., Jackson, G.D., Connelly, A., 2010. Track-density imaging (TDI): Super-resolution white matter imaging using whole-brain track-density mapping. Neuroimage. https://doi.org/10.1016/j.neuroimage.2010.07.024

Calzavara, R., Zappalà, A., Rozzi, S., Matelli, M., Luppino, G., 2005. Neurochemical characterization of the cerebellar-recipient motor thalamic territory in the macaque monkey. Eur. J. Neurosci. 21, 1869–1894. https://doi.org/10.1111/j.1460-9568.2005.04020.x

Chakravarty, M.M., Bertrand, G., Hodge, C.P., Sadikot, A.F., Collins, D.L., 2006. The creation of a brain atlas for image guided neurosurgery using serial histological data. Neuroimage. https://doi.org/10.1016/j.neuroimage.2005.09.041

Clayden, J.D., Thomas, D.L., Kraskov, A., 2019. Tractography-based parcellation does not provide strong evidence of anatomical organisation within the thalamus. Neuroimage 199, 418–426. https://doi.org/10.1016/j.neuroimage.2019.06.019

Crum, W.R., Camara, O., Hill, D.L.G., 2006. Generalized overlap measures for evaluation and validation in medical image analysis. IEEE Trans. Med. Imaging. https://doi.org/10.1109/TMI.2006.880587

Cury, R.G., Fraix, V., Castrioto, A., Pérez Fernández, M.A., Krack, P., Chabardes, S., Seigneuret, E., Alho, E.J.L., Benabid, A.-L., Moro, E., 2017. Thalamic deep brain stimulation for tremor in Parkinson disease, essential tremor, and dystonia. Neurology 89, 1416–1423. https://doi.org/10.1212/WNL.0000000000004295

da Silva, N.M., Ahmadi, S.-A., Tafula, S.N., Cunha, J.P.S., Bötzel, K., Vollmar, C., Rozanski, V.E., 2017. A diffusion-based connectivity map of the GPi for optimised stereotactic targeting in DBS. Neuroimage 144, 83–91. https://doi.org/10.1016/j.neuroimage.2016.06.018

Darian-Smith, C., Darian-Smith, I., Cheema, S.S., 1990. Thalamic projections to sensorimotor cortex in the macaque monkey: Use of multiple retrograde fluorescent tracers. J. Comp. Neurol. 299, 17–46. https://doi.org/10.1002/cne.902990103

Deuschl, G., Krack, P., Lauk, M., Timmer, J., 1996. Clinical neurophysiology of tremor. J. Clin. Neurophysiol. https://doi.org/10.1097/00004691-199603000-00002

Dhollander, T., Mito, R., Raffelt, D., Connelly, A., 2019. Improved white matter response function estimation for 3-tissue constrained spherical deconvolution. Proc. Intl. Soc. Mag. Reson. Med 555.

Dice, L.R., 1945. Measures of the Amount of Ecologic Association Between Species. Ecology. https://doi.org/10.2307/1932409

Domin, M., Lotze, M., 2019. Parcellation of motor cortex-associated regions in the human corpus callosum on the basis of Human Connectome Project data. Brain Struct. Funct. https://doi.org/10.1007/s00429-019-01849-1

Dum, R.P., Strick, P.L., 2003. An unfolded map of the cerebellar dentate nucleus and its projections to the cerebral cortex. J. Neurophysiol. 89, 634–639. https://doi.org/10.1152/jn.00626.2002

Eickhoff, S.B., Thirion, B., Varoquaux, G., Bzdok, D., 2015. Connectivity-based parcellation: Critique and implications. Hum. Brain Mapp. https://doi.org/10.1002/hbm.22933

Elias, G.J.B., Boutet, A., Joel, S.E., Germann, J., Gwun, D., Neudorfer, C., Gramer, R.M., Algarni, M., Paramanandam, V., Prasad, S., Beyn, M.E., Horn, A., Madhavan, R., Ranjan, M., Lozano, C.S., Kühn, A.A., Ashe, J., Kucharczyk, W., Munhoz, R.P., Giacobbe, P., Kennedy, S.H., Woodside, D.B., Kalia, S.K., Fasano, A., Hodaie, M., Lozano, A.M., 2020. Probabilistic Mapping of Deep Brain Stimulation: Insights from 15 Years of Therapy. Ann. Neurol. https://doi.org/10.1002/ana.25975

Elias, W.J., Huss, D., Voss, T., Loomba, J., Khaled, M., Zadicario, E., Frysinger, R.C., Sperling, S.A., Wylie, S., Monteith, S.J., Druzgal, J., Shah, B.B., Harrison, M., Wintermark, M., 2013. A pilot study of focused ultrasound thalamotomy for essential tremor. N. Engl. J. Med. 369, 640–648. https://doi.org/10.1056/NEJMoa1300962

Elias, W.J., Zheng, Z.A., Domer, P., Quigg, M., Pouratian, N., 2012. Validation of connectivity-based thalamic segmentation with direct electrophysiologic recordings from human sensory thalamus. Neuroimage 59, 2025–2034. https://doi.org/10.1016/j.neuroimage.2011.10.049

Essayed, W.I., Zhang, F., Unadkat, P., Cosgrove, G.R., Golby, A.J., O’Donnell, L.J., 2017. White matter tractography for neurosurgical planning: A topography-based review of the current state of the art. NeuroImage Clin. https://doi.org/10.1016/j.nicl.2017.06.011

Ewert, S., Plettig, P., Li, N., Chakravarty, M.M., Collins, D.L., Herrington, T.M., Kühn, A.A., Horn, A., 2018. Toward defining deep brain stimulation targets in MNI space: A subcortical atlas based on multimodal MRI, histology and structural connectivity. Neuroimage. https://doi.org/10.1016/j.neuroimage.2017.05.015

Fan, Y., Nickerson, L.D., Li, H., Ma, Y., Lyu, B., Miao, X., Zhuo, Y., Ge, J., Zou, Q., Gao, J.-H., 2015. Functional Connectivity-Based Parcellation of the Thalamus: An Unsupervised Clustering Method and Its Validity Investigation. Brain Connect. 5, 620–630. https://doi.org/10.1089/brain.2015.0338

Farquharson, S., Tournier, J.-D., Calamante, F., Fabinyi, G., Schneider-Kolsky, M., Jackson, G.D., Connelly, A., 2013. White matter fiber tractography: why we need to move beyond DTI. J. Neurosurg. https://doi.org/10.3171/2013.2.JNS121294

Fenoy, A.J., Schiess, M.C., 2018. Comparison of tractography-assisted to atlas-based targeting for deep brain stimulation in essential tremor. Mov. Disord. 33, 1895–1901. https://doi.org/10.1002/mds.27463

Fenoy, A.J., Schiess, M.C., 2017. Deep Brain Stimulation of the Dentato-Rubro-Thalamic Tract: Outcomes of Direct Targeting for Tremor. Neuromodulation Technol. Neural Interface 20, 429–436. https://doi.org/10.1111/ner.12585

Fiechter, M., Nowacki, A., Oertel, M.F., Fichtner, J., Debove, I., Lachenmayer, M.L., Wiest, R., Bassetti, C.L., Raabe, A., Kaelin-Lang, A., Schüpbach, M.W., Pollo, C., 2017. Deep Brain Stimulation for Tremor: Is There a Common Structure? Stereotact. Funct. Neurosurg. 95, 243– 250. https://doi.org/10.1159/000478270

Fonov, V., Evans, A., McKinstry, R., Almli, C., Collins, D., 2009. Unbiased nonlinear average age-appropriate brain templates from birth to adulthood. Neuroimage. https://doi.org/10.1016/s1053-8119(09)70884-5

Gallay, M.N., Jeanmonod, D., Liu, J., Morel, A., 2008. Human pallidothalamic and cerebellothalamic tracts: Anatomical basis for functional stereotactic neurosurgery. Brain Struct. Funct. https://doi.org/10.1007/s00429-007-0170-0

Girard, G., Caminiti, R., Battaglia-Mayer, A., St-Onge, E., Ambrosen, K.S., Eskildsen, S.F., Krug, K., Dyrby, T.B., Descoteaux, M., Thiran, J.P., Innocenti, G.M., 2020. On the cortical connectivity in the macaque brain: A comparison of diffusion tractography and histological tracing data. Neuroimage 221, 117201. https://doi.org/10.1016/j.neuroimage.2020.117201

Glasser, M.F., Smith, S.M., Marcus, D.S., Andersson, J.L.R., Auerbach, E.J., Behrens, T.E.J., Coalson, T.S., Harms, M.P., Jenkinson, M., Moeller, S., Robinson, E.C., Sotiropoulos, S.N., Xu, J., Yacoub, E., Ugurbil, K., Van Essen, D.C., 2016. The Human Connectome Project’s neuroimaging approach. Nat. Neurosci. 19, 1175–1187. https://doi.org/10.1038/nn.4361

Glasser, M.F., Sotiropoulos, S.N., Wilson, J.A., Coalson, T.S., Fischl, B., Andersson, J.L., Xu, J., Jbabdi, S., Webster, M., Polimeni, J.R., Van Essen, D.C., Jenkinson, M., 2013. The minimal preprocessing pipelines for the Human Connectome Project. Neuroimage 80, 105–124. https://doi.org/10.1016/j.neuroimage.2013.04.127

Hassler, 1983. Stereotaxy of the human brain — anatomical, physiological and clinical applications. Clin. Neurol. Neurosurg. https://doi.org/10.1016/0303-8467(83)90036-7

Helmich, R.C., Toni, I., Deuschl, G., Bloem, B.R., 2013. The pathophysiology of essential tremor and parkinson’s tremor. Curr. Neurol. Neurosci. Rep. https://doi.org/10.1007/s11910-013-0378-8

Horn, A., Kühn, A.A., Merkl, A., Shih, L., Alterman, R., Fox, M., 2017. Probabilistic conversion of neurosurgical DBS electrode coordinates into MNI space. Neuroimage. https://doi.org/10.1016/j.neuroimage.2017.02.004

Iglesias, J.E., Insausti, R., Lerma-Usabiaga, G., Bocchetta, M., Van Leemput, K., Greve, D.N., van der Kouwe, A., Fischl, B., Caballero-Gaudes, C., Paz-Alonso, P.M., 2018. A probabilistic atlas of the human thalamic nuclei combining ex vivo MRI and histology. Neuroimage 183, 314– 326. https://doi.org/10.1016/j.neuroimage.2018.08.012

Ilinsky, I., Horn, A., Paul-Gilloteaux, P., Gressens, P., Verney, C., Kultas-Ilinsky, K., 2018. Human Motor Thalamus Reconstructed in 3D from Continuous Sagittal Sections with Identified Subcortical Afferent Territories. eneuro 5, ENEURO.0060-18.2018. https://doi.org/10.1523/ENEURO.0060-18.2018

Jbabdi, S., Johansen-Berg, H., 2011. Tractography: Where Do We Go from Here? Brain Connect. https://doi.org/10.1089/brain.2011.0033

Jeurissen, B., Descoteaux, M., Mori, S., Leemans, A., 2019. Diffusion MRI fiber tractography of the brain. NMR Biomed. https://doi.org/10.1002/nbm.3785

Jeurissen, B., Tournier, J.D., Dhollander, T., Connelly, A., Sijbers, J., 2014. Multi-tissue constrained spherical deconvolution for improved analysis of multi-shell diffusion MRI data. Neuroimage. https://doi.org/10.1016/j.neuroimage.2014.07.061

Johansen-Berg, H., Behrens, T.E.J., Sillery, E., Ciccarelli, O., Thompson, A.J., Smith, S.M., Matthews, P.M., 2005. Functional-anatomical validation and individual variation of diffusion tractography-based segmentation of the human thalamus. Cereb. Cortex 15, 31–39. https://doi.org/10.1093/cercor/bhh105

Jones, D.K., Horsfield, M.A., Simmons, A., 1999. Optimal strategies for measuring diffusion in anisotropic systems by magnetic resonance imaging. Magn. Reson. Med. 42, 515–525. https://doi.org/10.1002/(SICI)1522-2594(199909)42:3<515::AID-MRM14>3.0.CO;2-Q

Kincses, Z.T., Szabó, N., Valálik, I., Kopniczky, Z., Dézsi, L., Klivényi, P., Jenkinson, M., Király, A., Babos, M., Vörös, E., Barzó, P., Vécsei, L., 2012. Target identification for stereotactic thalamotomy using diffusion tractography. PLoS One 7. https://doi.org/10.1371/journal.pone.0029969

Krishna, V., Sammartino, F., Agrawal, P., Changizi, B.K., Bourekas, E., Knopp, M. V., Rezai, A., 2019. Prospective tractography-based targeting for improved safety of focused ultrasound thalamotomy. Clin. Neurosurg. https://doi.org/10.1093/neuros/nyy020

Lambert, C., Simon, H., Colman, J., Barrick, T.R., 2017. Defining thalamic nuclei and topographic connectivity gradients in vivo. Neuroimage 158, 466–479. https://doi.org/10.1016/j.neuroimage.2016.08.028

Middlebrooks, E.H., Domingo, R.A., Vivas-Buitrago, T., Okromelidze, L., Tsuboi, T., Wong, J.K., Eisinger, R.S., Almeida, L., Burns, M.R., Horn, A., Uitti, R.J., Wharen, R.E., Holanda, V.M., Grewal, S.S., 2020. Neuroimaging advances in deep brain stimulation: Review of indications, anatomy, and brain connectomics. Am. J. Neuroradiol. https://doi.org/10.3174/ajnr.A6693

Middlebrooks, E.H., Holanda, V.M., Tuna, I.S., Deshpande, H.D., Bredel, M., Almeida, L., Walker, H.C., Guthrie, B.L., Foote, K.D., Okun, M.S., 2018a. A method for pre-operative single-subject thalamic segmentation based on probabilistic tractography for essential tremor deep brain stimulation. Neuroradiology. https://doi.org/10.1007/s00234-017-1972-2

Middlebrooks, E.H., Tuna, I.S., Almeida, L., Grewal, S.S., Wong, J., Heckman, M.G., Lesser, E.R., Bredel, M., Foote, K.D., Okun, M.S., Holanda, V.M., 2018b. Structural connectivity–based segmentation of the thalamus and prediction of tremor improvement following thalamic deep brain stimulation of the ventral intermediate nucleus. NeuroImage Clin. 20, 1266–1273. https://doi.org/10.1016/j.nicl.2018.10.009

Middlebrooks, E.H., Tuna, I.S., Grewal Grewal, S.S., Almeida, L. eonardo. Heckman, M.G., Lesser, E.R., Foote, K.D., Okun, M.S., Holanda, V.M., 2018c. Segmentation of the globus pallidus internus using probabilistic diffusion tractography for deep brain stimulation targeting in Parkinson disease. Am. J. Neuroradiol. 39, 1127–1134. https://doi.org/10.3174/ajnr.A5641

Miller, T.R., Zhuo, J., Eisenberg, H.M., Fishman, P.S., Melhem, E.R., Gullapalli, R., Gandhi, D., 2019. Targeting of the dentato-rubro-thalamic tract for MR-guided focused ultrasound treatment of essential tremor. Neuroradiol. J. 32, 401–407. https://doi.org/10.1177/1971400919870180

Morel, A., Magnin, M., Jeanmonod, D., 1997. Multiarchitectonic and stereotactic atlas of the human thalamus. J. Comp. Neurol. 387, 588–630. https://doi.org/10.1002/(SICI)1096-9861(19971103)387:4<588::AID-CNE8>3.0.CO;2-Z

O’muircheartaigh, J., Keller, S.S., Barker, G.J., Richardson, M.P., 2015. White matter connectivity of the thalamus delineates the functional architecture of competing thalamocortical systems. Cereb. Cortex 25, 4477–4489. https://doi.org/10.1093/cercor/bhv063

Palesi, F., Tournier, J.D., Calamante, F., Muhlert, N., Castellazzi, G., Chard, D., D’Angelo, E., Wheeler-Kingshott, C.A.M., 2015. Contralateral cerebello-thalamo-cortical pathways with prominent involvement of associative areas in humans in vivo. Brain Struct. Funct. https://doi.org/10.1007/s00429-014-0861-2

Palesi, F., Tournier, J.D., Calamante, F., Muhlert, N., Castellazzi, G., Chard, D., D’Angelo, E., Wheeler-Kingshott, C.G., 2016. Reconstructing contralateral fiber tracts: Methodological aspects of cerebello-thalamo-cortical pathway reconstruction. Funct. Neurol. https://doi.org/10.11138/FNeur/2016.31.4.229

Papavassiliou, E., Rau, G., Heath, S., Abosch, A., Barbaro, N.M., Larson, P.S., Lamborn, K., Starr, P.A., Sharan, A.D., Rezai, A.R., Benabid, A.L., Lozano, A.M., 2004. Thalamic Deep Brain Stimulation for Essential Tremor: Relation of Lead Location to Outcome. Neurosurgery. https://doi.org/10.1227/01.NEU.0000119329.66931.9E

Patenaude, B., Smith, S.M., Kennedy, D.N., Jenkinson, M., 2011. A Bayesian model of shape and appearance for subcortical brain segmentation. Neuroimage 56, 907–922. https://doi.org/10.1016/j.neuroimage.2011.02.046

Patriat, R., Cooper, S.E., Duchin, Y., Niederer, J., Lenglet, C., Aman, J., Park, M.C., Vitek, J.L., Harel, N., 2018. Individualized tractography-based parcellation of the globus pallidus pars interna using 7T MRI in movement disorder patients prior to DBS surgery. Neuroimage 178, 198–209. https://doi.org/10.1016/j.neuroimage.2018.05.048

Petersen, M. V., Lund, T.E., Sunde, N., Frandsen, J., Rosendal, F., Juul, N., Østergaard, K., 2017. Probabilistic versus deterministic tractography for delineation of the cortico-subthalamic hyperdirect pathway in patients with Parkinson disease selected for deep brain stimulation. J. Neurosurg. https://doi.org/10.3171/2016.4.JNS1624

Plantinga, B.R., Temel, Y., Duchin, Y., Uludağ, K., Patriat, R., Roebroeck, A., Kuijf, M., Jahanshahi, A., ter Haar Romenij, B., Vitek, J., Harel, N., 2018. Individualized parcellation of the subthalamic nucleus in patients with Parkinson’s disease with 7T MRI. Neuroimage. https://doi.org/10.1016/j.neuroimage.2016.09.023

Pouratian, N., Zheng, Z., Bari, A.A., Behnke, E., Elias, W.J., DeSalles, A.A.F., 2011. Multi-institutional evaluation of deep brain stimulation targeting using probabilistic connectivity-based thalamic segmentation: Clinical article. J. Neurosurg. 115, 995–1004. https://doi.org/10.3171/2011.7.JNS11250

Reuter, N., Genon, S., Kharabian Masouleh, S., Hoffstaedter, F., Liu, X., Kalenscher, T., Eickhoff, S.B., Patil, K.R., 2020. CBPtools: a Python package for regional connectivity-based parcellation. Brain Struct. Funct. https://doi.org/10.1007/s00429-020-02046-1

Sammartino, F., Krishna, V., King, N.K.K., Lozano, A.M., Schwartz, M.L., Huang, Y., Hodaie, M., 2016. Tractography-Based Ventral Intermediate Nucleus Targeting: Novel Methodology and Intraoperative Validation. Mov. Disord. https://doi.org/10.1002/mds.26633

Saygin, Z.M., Osher, D.E., Augustinack, J., Fischl, B., Gabrieli, J.D.E., 2011. Connectivity-based segmentation of human amygdala nuclei using probabilistic tractography. Neuroimage. https://doi.org/10.1016/j.neuroimage.2011.03.006

Schlaier, J.R., Beer, A.L., Faltermeier, R., Fellner, C., Steib, K., Lange, M., Greenlee, M.W., Brawanski, A.T., Anthofer, J.M., 2017. Probabilistic vs. deterministic fiber tracking and the influence of different seed regions to delineate cerebellar-thalamic fibers in deep brain stimulation. Eur. J. Neurosci. https://doi.org/10.1111/ejn.13575

Schnitzler, A., Münks, C., Butz, M., Timmermann, L., Gross, J., 2009. Synchronized brain network associated with essential tremor as revealed by magnetoencephalography. Mov. Disord. https://doi.org/10.1002/mds.22633

Shah, B.R., Lehman, V.T., Kaufmann, T.J., Blezek, D., Waugh, J., Imphean, D., Yu, F.F., Patel, T.R., Chitnis, S., Dewey, R.B., Maldjian, J.A., Chopra, R., 2020. Advanced MRI techniques for transcranial high intensity focused ultrasound targeting. Brain 143, 2664–2672. https://doi.org/10.1093/brain/awaa107

Sinke, M.R.T., Otte, W.M., Christiaens, D., Schmitt, O., Leemans, A., van der Toorn, A., Sarabdjitsingh, R.A., Joëls, M., Dijkhuizen, R.M., 2018. Diffusion MRI-based cortical connectome reconstruction: dependency on tractography procedures and neuroanatomical characteristics. Brain Struct. Funct. 223, 2269–2285. https://doi.org/10.1007/s00429-018-1628-y

Smith, S.M., 2002. Fast robust automated brain extraction. Hum. Brain Mapp. 17, 143–155. https://doi.org/10.1002/hbm.10062

Smith, S.M., Jenkinson, M., Woolrich, M.W., Beckmann, C.F., Behrens, T.E.J., Johansen-Berg, H., Bannister, P.R., De Luca, M., Drobnjak, I., Flitney, D.E., Niazy, R.K., Saunders, J., Vickers, J., Zhang, Y., De Stefano, N., Brady, J.M., Matthews, P.M., 2004. Advances in functional and structural MR image analysis and implementation as FSL, in: NeuroImage. https://doi.org/10.1016/j.neuroimage.2004.07.051

Sotiropoulos, S.N., Jbabdi, S., Xu, J., Andersson, J.L., Moeller, S., Auerbach, E.J., Glasser, M.F., Hernandez, M., Sapiro, G., Jenkinson, M., Feinberg, D.A., Yacoub, E., Lenglet, C., Van Essen, D.C., Ugurbil, K., Behrens, T.E.J., 2013. Advances in diffusion MRI acquisition and processing in the Human Connectome Project. Neuroimage. https://doi.org/10.1016/j.neuroimage.2013.05.057

Su, J.H., Thomas, F.T., Kasoff, W.S., Tourdias, T., Choi, E.Y., Rutt, B.K., Saranathan, M., 2019. Thalamus Optimized Multi Atlas Segmentation (THOMAS): fast, fully automated segmentation of thalamic nuclei from structural MRI. Neuroimage. https://doi.org/10.1016/j.neuroimage.2019.03.021

Sudhyadhom, A., McGregor, K., Okun, M.S., Foote, K.D., Trinastic, J., Crosson, B., Bova, F.J., 2013. Delineation of motor and somatosensory thalamic subregions utilizing probabilistic diffusion tractography and electrophysiology. J. Magn. Reson. Imaging 37, 600–609. https://doi.org/10.1002/jmri.23861

Tang, Y., Sun, W., Toga, A.W., Ringman, J.M., Shi, Y., 2018. A probabilistic atlas of human brainstem pathways based on connectome imaging data. Neuroimage 169, 227–239. https://doi.org/10.1016/j.neuroimage.2017.12.042

Theisen, F., Leda, R., Pozorski, V., Oh, J.M., Adluru, N., Wong, R., Okonkwo, O., Dean, D.C., Bendlin, B.B., Johnson, S.C., Alexander, A.L., Gallagher, C.L., 2017. Evaluation of striatonigral connectivity using probabilistic tractography in Parkinson’s disease. NeuroImage Clin. https://doi.org/10.1016/j.nicl.2017.09.009

Tian, Q., Wintermark, M., Jeffrey Elias, W., Ghanouni, P., Halpern, C.H., Henderson, J.M., Huss, D.S., Goubran, M., Thaler, C., Airan, R., Zeineh, M., Pauly, K.B., McNab, J.A., 2018. Diffusion MRI tractography for improved transcranial MRI-guided focused ultrasound thalamotomy targeting for essential tremor. NeuroImage Clin. 19, 572–580. https://doi.org/10.1016/j.nicl.2018.05.010

Tournier, J.D., Calamante, F., Connelly, A., 2007. Robust determination of the fibre orientation distribution in diffusion MRI: Non-negativity constrained super-resolved spherical deconvolution. Neuroimage. https://doi.org/10.1016/j.neuroimage.2007.02.016

Tournier, J.D., Smith, R., Raffelt, D., Tabbara, R., Dhollander, T., Pietsch, M., Christiaens, D., Jeurissen, B., Yeh, C.H., Connelly, A., 2019. MRtrix3: A fast, flexible and open software framework for medical image processing and visualisation. Neuroimage. https://doi.org/10.1016/j.neuroimage.2019.116137

Tournier, J.D., Yeh, C.H., Calamante, F., Cho, K.H., Connelly, A., Lin, C.P., 2008. Resolving crossing fibres using constrained spherical deconvolution: Validation using diffusion-weighted imaging phantom data. Neuroimage 42, 617–625. https://doi.org/10.1016/j.neuroimage.2008.05.002

Traynor, C., Heckemann, R.A., Hammers, A., O’Muircheartaigh, J., Crum, W.R., Barker, G.J., Richardson, M.P., 2010. Reproducibility of thalamic segmentation based on probabilistic tractography. Neuroimage 52, 69–85. https://doi.org/10.1016/j.neuroimage.2010.04.024

Tsolaki, E., Downes, A., Speier, W., Elias, W.J., Pouratian, N., 2018. The potential value of probabilistic tractography-based for MR-guided focused ultrasound thalamotomy for essential tremor. NeuroImage Clin. 17, 1019–1027. https://doi.org/10.1016/j.nicl.2017.12.018

Van Essen, D.C., 1997. A tension-based theory of morphogenesis and compact wiring in the central nervous system. Nature. https://doi.org/10.1038/385313a0

Van Essen, D.C., Smith, S.M., Barch, D.M., Behrens, T.E.J., Yacoub, E., Ugurbil, K., 2013. The WU-Minn Human Connectome Project: An overview. Neuroimage 80, 62–79. https://doi.org/10.1016/j.neuroimage.2013.05.041

Van Essen, D.C., Ugurbil, K., Auerbach, E., Barch, D., Behrens, T.E.J., Bucholz, R., Chang, A., Chen, L., Corbetta, M., Curtiss, S.W., Della Penna, S., Feinberg, D., Glasser, M.F., Harel, N., Heath, A.C., Larson-Prior, L., Marcus, D., Michalareas, G., Moeller, S., Oostenveld, R., Petersen, S.E., Prior, F., Schlaggar, B.L., Smith, S.M., Snyder, A.Z., Xu, J., Yacoub, E., 2012. The Human Connectome Project: A data acquisition perspective. Neuroimage 62, 2222–2231. https://doi.org/10.1016/j.neuroimage.2012.02.018

Wasserthal, J., Neher, P., Maier-Hein, K.H., 2018. TractSeg - Fast and accurate white matter tract segmentation. Neuroimage 183, 239–253. https://doi.org/10.1016/j.neuroimage.2018.07.070

Zhang, D., Snyder, A.Z., Fox, M.D., Sansbury, M.W., Shimony, J.S., Raichle, M.E., 2008. Intrinsic Functional Relations Between Human Cerebral Cortex and Thalamus. J. Neurophysiol. 100, 1740–1748. https://doi.org/10.1152/jn.90463.2008

