## Supplementary Information for "Ventral Intermediate Nucleus structural connectivity-derived segmentation: anatomical reliability and variability"

#### *Supplementary methods - Region of Interest (ROI) selection*

Regions of interest (ROI) employed to perform tractography-based parcellation of thalamus are depicted in Supplementary Figure 1. Masks of cortical targets were retrieved from the automated anatomical labeling atlas (AAL3) (Rolls et al., 2020). Specifically, the atlas was registered from standard space to each subject native space using non-linear transformations, and the following cortical ROIs were extracted: precentral gyrus, supplementary motor area (SMA), postcentral gyrus, paracentral lobule, prefrontal (by merging superior, middle, inferior frontal gyri, anterior and middle cingulate cortices and orbitofrontal cortex ROIs), parietal (superior and inferior parietal lobules, supramarginal gyrus, angular gyrus, precuneus), temporal (Heschl gyrus, temporal pole, superior, middle and inferior temporal gyri) and occipital (calcarine cortex, cuneus, lingual gyrus, fusiform gyrus, superior, middle and inferior occipital gyri) cortices (Behrens et al., 2003; Johansen-Berg et al., 2005).

Finally, the ROIs (superior cerebellar peduncle and red nucleus) needed to reconstruct the DRTC tract were obtained applying a manual segmentation protocol on track-density images (TDI), similarly to what has been performed in (Tang et al., 2018). Specifically, whole-brain tractography with default tracking parameters has been performed on each subject in order to obtain a 10 million streamlines tractogram. TDI maps have been obtained by sampling each whole brain tractogram to maps within a 0.7 mm resolution grid, which allows direct visualization of anatomical structures not easily identified on conventional MRI (Calamante, 2016; Calamante et al., 2012, 2010).  The masks of left and right superior cerebellar peduncles (SCPs) were defined on a single coronal slice for each subject. Finally, the red nucleus have been obtained from DISTAL atlas (Ewert et al., 2018), then superimposed on TDI maps and manually corrected, as it appears clearly distinguishable as a round hypointense structure in axial sections. The entire segmentation process has been carried out by a trained neuroanatomist (D.M).

#### References


Behrens, T.E.J., Woolrich, M.W., Smith, S.M., Boulby, P.A., Barker, G.J., Sillery, E.L., Sheehan, K., Ciccarelli, O., Thompson, A.J., Brady, J.M., Matthews, P.M., 2003. Non-invasive mapping of connecitons between human thalamus and cortex using DTI. Nat. Neurosci.

Calamante, F., 2016. Super-resolution track density imaging: Anatomic detail versus quantification. Am. J. Neuroradiol. https://doi.org/10.3174/ajnr.A4721

Calamante, F., Tournier, J.D., Jackson, G.D., Connelly, A., 2010. Track-density imaging (TDI): Super-resolution white matter imaging using whole-brain track-density mapping. Neuroimage. https://doi.org/10.1016/j.neuroimage.2010.07.024

Calamante, F., Tournier, J.D., Smith, R.E., Connelly, A., 2012. A generalised framework for super-resolution track-weighted imaging. Neuroimage. https://doi.org/10.1016/j.neuroimage.2011.08.099

Ewert, S., Plettig, P., Li, N., Chakravarty, M.M., Collins, D.L., Herrington, T.M., Kühn, A.A., Horn, A., 2018. Toward defining deep brain stimulation targets in MNI space: A subcortical atlas based on multimodal MRI, histology and structural connectivity. Neuroimage 170, 271–282. https://doi.org/10.1016/j.neuroimage.2017.05.015

Johansen-Berg, H., Behrens, T.E.J., Sillery, E., Ciccarelli, O., Thompson, A.J., Smith, S.M., Matthews, P.M., 2005. Functional-anatomical validation and individual variation of diffusion tractography-based segmentation of the human thalamus. Cereb. Cortex 15, 31–39. https://doi.org/10.1093/cercor/bhh105

Rolls, E.T., Huang, C.-C., Lin, C.-P., Feng, J., Joliot, M., 2020. Automated anatomical labelling atlas 3. Neuroimage 206, 116189. https://doi.org/10.1016/j.neuroimage.2019.116189

Tang, Y., Sun, W., Toga, A.W., Ringman, J.M., Shi, Y., 2018. A probabilistic atlas of human brainstem pathways based on connectome imaging data. Neuroimage 169, 227–239. https://doi.org/10.1016/j.neuroimage.2017.12.042

*Supplementary Figures*

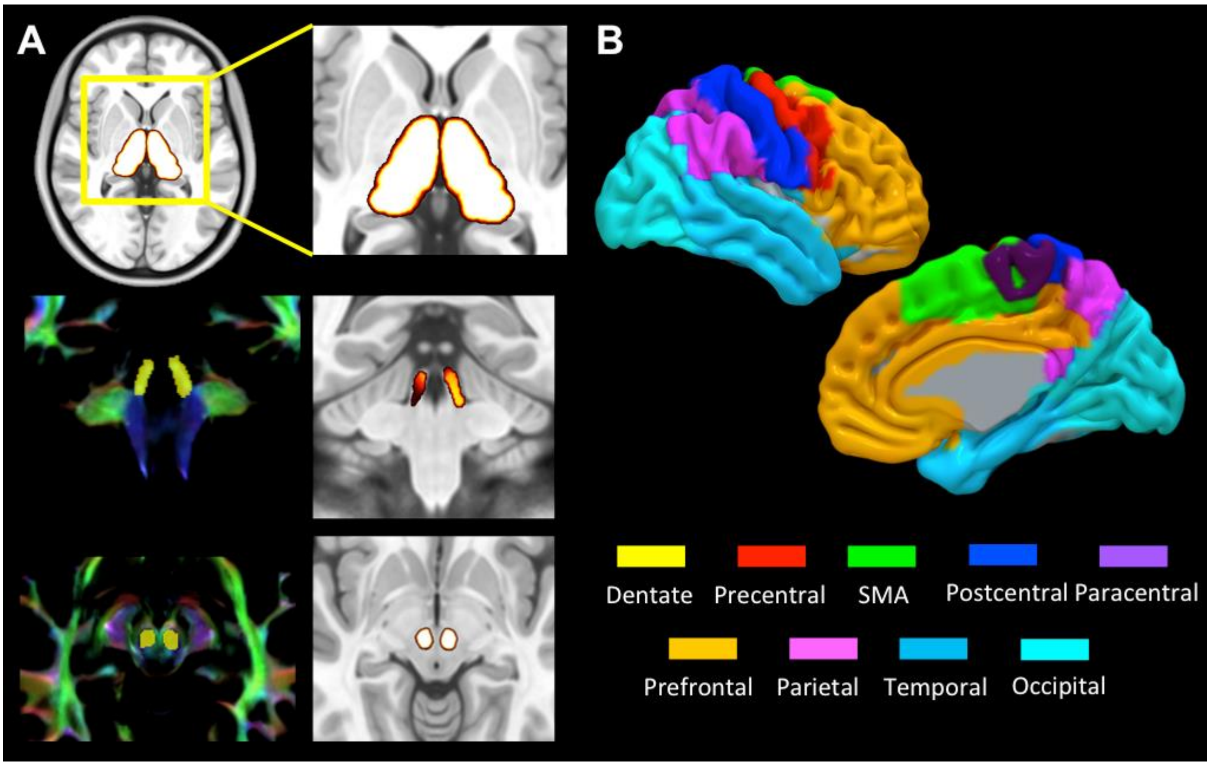


**Supplementary Figure 1. ROI selection**. Subcortical and cortical regions of interest employed to perform connectivity-based parcellation. (A) The upper row depicts the thalamus as obtained from FSL FIRST tool. The mid and bottom rows show respectively the ROI of the superior cerebellar peduncle and red nucleus as obtained applying manual segmentation on TDI maps. (B) Cortical areas retrieved from AAL3 atlas. Target ROI have been labeled according to the color-coding provided by the legend.

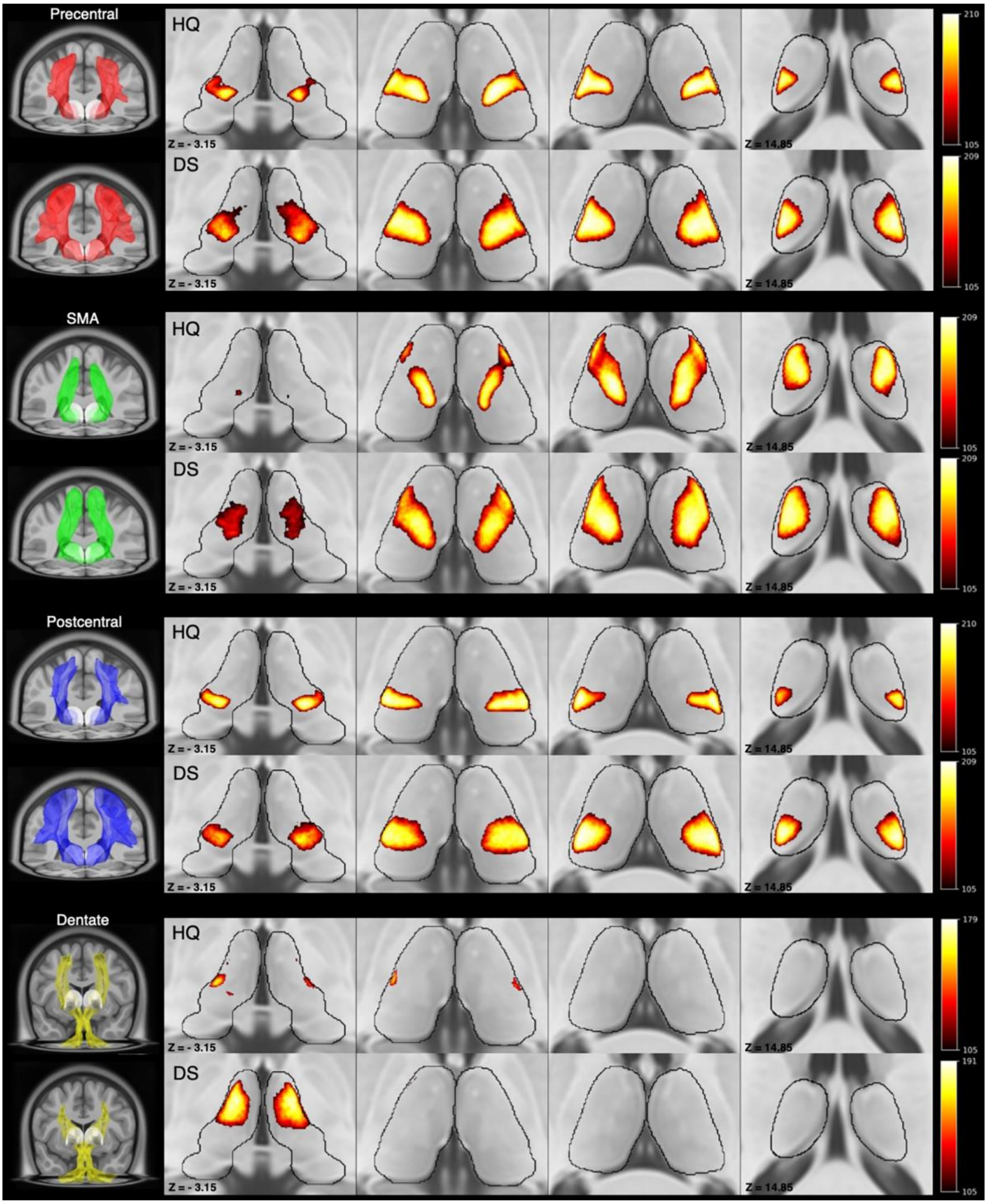


**Supplementary Figure 1. Thalamic MPMs.** Maximum Probability Maps (MPMs) of precentral, SMA, postcentral and dentate connectivity cluster are depicted in multiple axial slices, covering the entire thalamic volume. Each MPM is accompanied by a color bar where the lowest intensity value corresponds to the 50% threshold (-thr 105), whilst the highest intensity represents the number of subjects across which the depicted voxels overlapped. The most left column shows 3D volume rendering of population-level tracts: precentral (red), SMA (green), postcentral (blue), dentate (yellow).

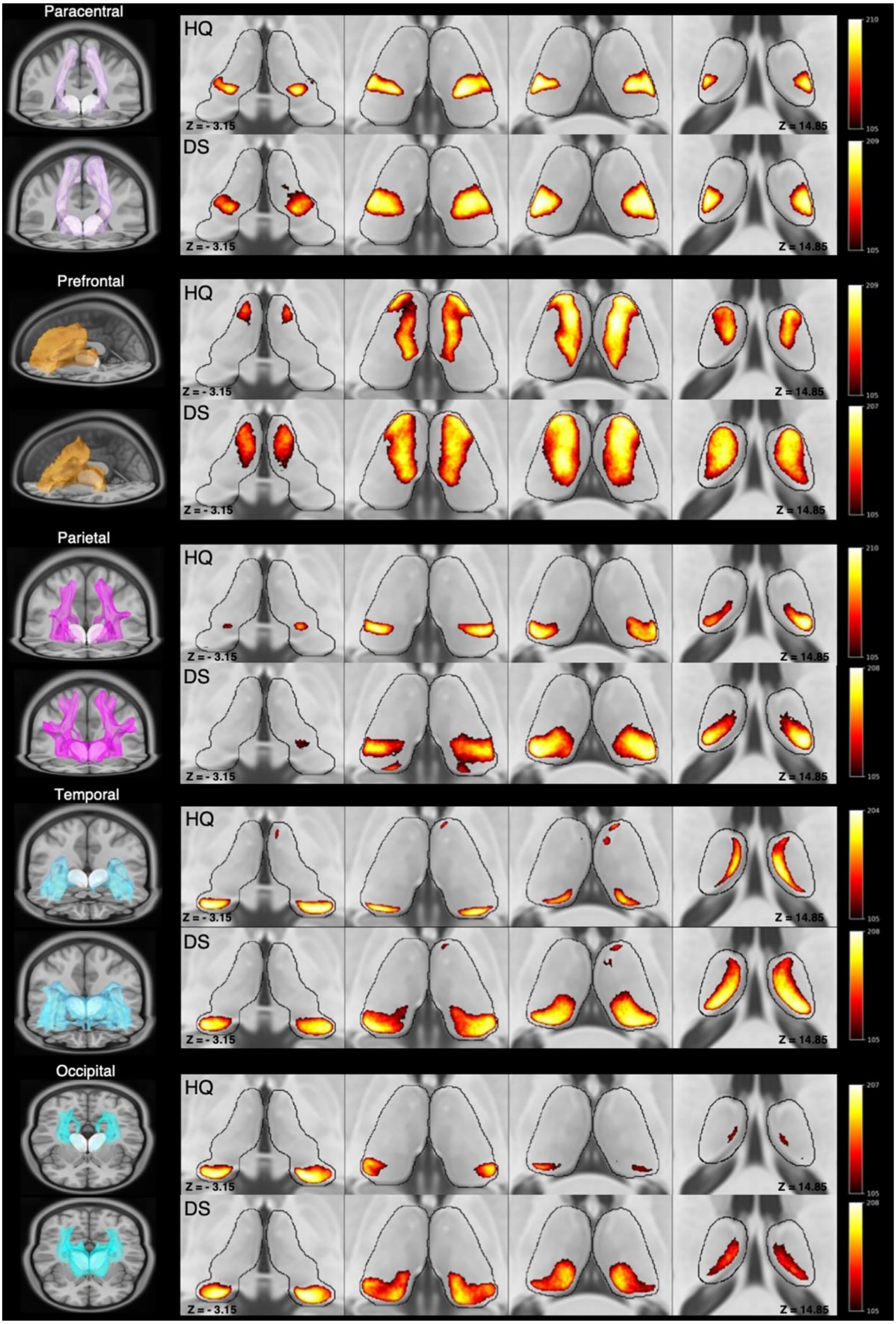


**Supplementary Figure 2. Thalamic MPMs.** Heat maps of paracentral, prefrontal, parietal, occipital, temporal MPMs are depicted in multiple axial slices extending from ventral to dorsal thalamus. Each MPM is accompanied by a color bar where the lowest intensity value corresponds to the 50% threshold (-thr 105) whilst the highest intensity represents the maximum number of subjects across which the depicted voxels overlapped. 3D volume rendering of tracts MPM corresponding to each cluster: paracentral (pink), prefrontal (orange), parietal (purple), temporal (light blue), occipital (cyan) are displayed on the extreme left.

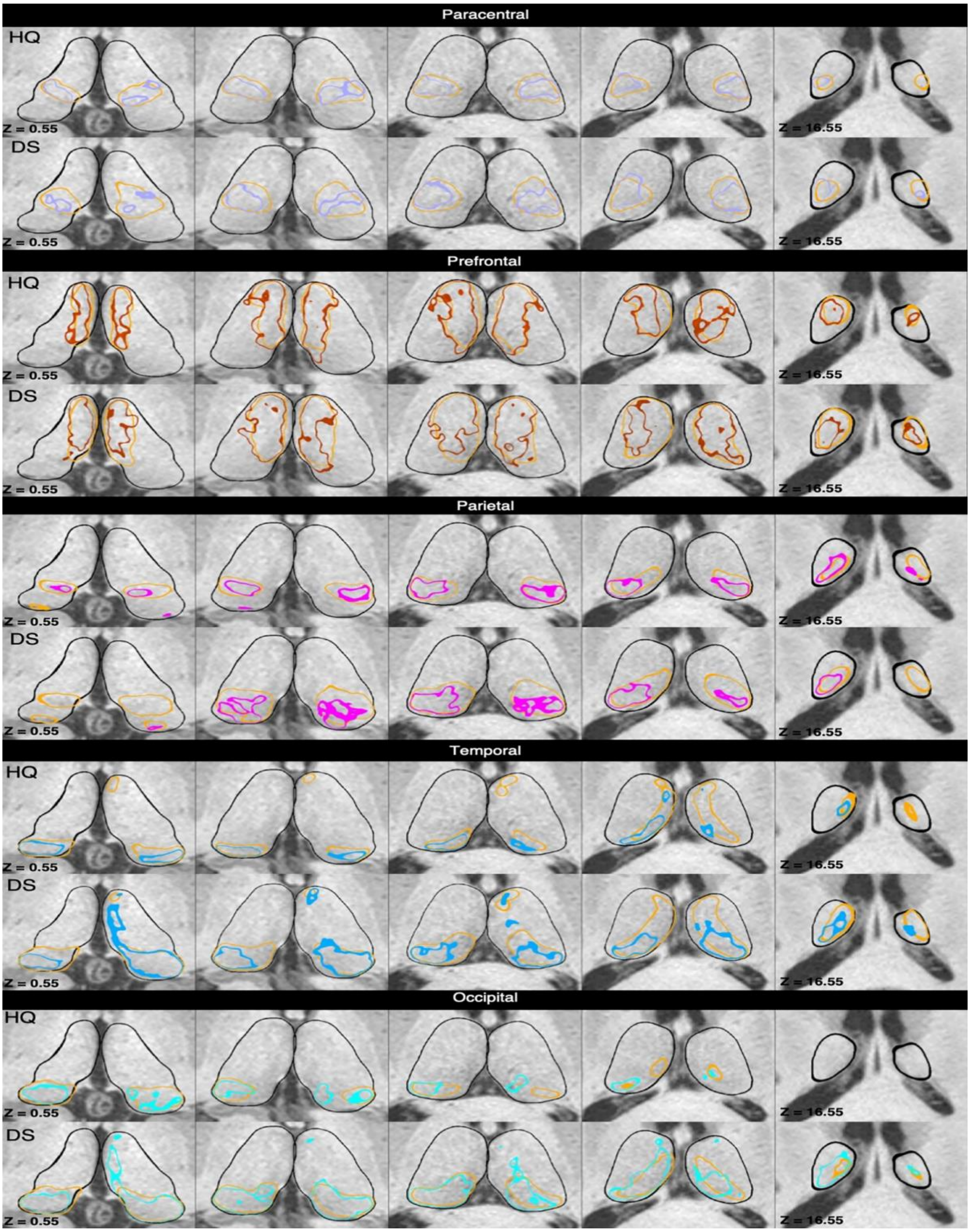


**Supplementary Figure 3. Comparison between tractographic atlas-based and personalized procedures.** This figure depicts the spatial relations between MPMs registered on a random subject (100408 from the 100 unrelated dataset) and the corresponding individualized maps. Multiple axial slices centered on the thalamic region show, from ventral to dorsal slices, boundaries of the MPM registered on subject native space (light orange) and those of the corresponding connectivity maps of paracentral (pink), prefrontal (orange), parietal (purple), temporal (light blue), occipital (cyan). For each cluster, the upper rows show the connectivity maps obtained from high quality data while the lower rows display those obtained from downsampled datasets.

### *Supplementary Tables*

| Clusters | High Quality | | | | Downsampled | | | |
|---|---|---|---|---|---|---|---|---|
| | Volume | COG ( x, y, z ) | | | Volume | COG ( x, y, z ) | | |
| **Left paracentral** | 12859 | -15.99 | -21.18 | 6.95 | 19994 | -15.76 | -20.61 | 7.19 |
| **Right paracentral** | 9238 | 16.54 | -21.02 | 6.38 | 15610 | 16.20 | -20.52 | 7.11 |
| **Left prefrontal** | 25913 | -9.64 | -12.80 | 7.95 | 38365 | -10.37 | -14.03 | 8.09 |
| **Right prefrontal** | 26525 | 8.76 | -12.33 | 7.86 | 38533 | 10.07 | -13.36 | 8.08 |
| **Left parietal** | 11247 | -17.11 | -25.28 | 9.09 | 22289 | -15.73 | -25.14 | 8.66 |
| **Right parietal** | 8242 | 17.02 | -25.41 | 8.28 | 17939 | 14.89 | -24.92 | 8.96 |
| **Left temporal** | 10936 | -12.86 | -24.33 | 7.89 | 27060 | -14.13 | -24.72 | 8.40 |
| **Right temporal** | 7211 | 13.53 | -26.69 | 7.12 | 20881 | 14.12 | -25.51 | 8.24 |
| **Left occipital** | 6819 | -17.88 | -30.47 | 2.35 | 19588 | -14.18 | -28.09 | 6.29 |
| **Right occipital** | 6862 | 17.60 | -29.31 | 3.23 | 18848 | 13.81 | -26.87 | 6.52 |

**Supplementary Table 1.** Volumes (in voxels) and COG (x, y, z) of paracentral, prefrontal, parietal, temporal and occipital MPMs obtained from the most reliable pipeline (CSD-THR), for high quality and downsampled dataset.

| Clusters | High Quality | | | | Downsampled | | | |
|---|---|---|---|---|---|---|---|---|
| | Avg DICE | | Avg Euclidean Distance | | Avg DICE | | Avg Euclidean Distance | |
| | Left | Right | Left | Right | Left | Right | Left | Right |
| **Paracentral** | 0.67 | 0.66 | 1.46 | 1.36 | 0.64 | 0.64 | 1.84 | 1.85 |
| **Prefrontal** | 0.71 | 0.68 | 1.41 | 1.61 | 0.70 | 0.69 | 1.66 | 1.88 |
| **Parietal** | 0.55 | 0.51 | 2.83 | 2.92 | 0.54 | 0.52 | 4.05 | 3.84 |
| **Temporal** | 0.52 | 0.45 | 2.67 | 4.34 | 0.61 | 0.58 | 3.34 | 4.27 |
| **Occipital** | 0.45 | 0.48 | 5.55 | 4.81 | 0.56 | 0.56 | 4.96 | 4.59 |

**Supplementary Table 2**. Average Dice coefficents and Euclidean distances between paracentral, prefrontal, parietal, temporal and occipital group-level and individualized maps in high quality and downsampled datasets. Both MPMs and individual maps come from the most reliable pipeline (CSD-THR).

| Avg Euclidean Distance | High Quality | | Downsampled | |
|---|---|---|---|---|
| | Left | Right | Left | Right |
| **Paracentral** | 2.78 | 2.53 | 1.29 | 1.32 |
| **Prefrontal** | 1.87 | 2.14 | 0.90 | 0.90 |
| **Parietal** | 2.99 | 2.92 | 1.57 | 1.65 |
| **Temporal** | 1.81 | 2.09 | 1.44 | 1.59 |
| **Occipital** | 4.06 | 4.22 | 1.72 | 1.73 |

**Supplementary Table 3.** Table summarizing average Euclidean distances between mean COG and individualized COG of paracentral, prefrontal, parietal, temporal and occipital connectivity clusters on the ICBM 2009b asymmetric template for both high quality and downsampled datasets.

*Effects of pipelines and data quality on connectivity clusters*

| CLUSTERS | SOURCE | TYPE III SUM OF SQUARES | DF* | ERROR | MEAN SQUARE | F | SIG. | $\eta_p^2$ |
|---|---|---|---|---|---|---|---|---|
| **L_Paracentral** | Dataquality | 7489.97 | 1 | 209 | 7489.97 | 2083.28 | .001 | .91 |
| | Pipeline | 159593.49 | 1.60 | 333.84 | 53197.83 | 5541.96 | .001 | .9 |
| | Dataquality * Pipeline | 13237.46 | 1.22 | 254.41 | 4412.49 | 1305.50 | .001 | .86 |
| **R_Paracentral** | Dataquality | 6590.36 | 1 | 209 | 6590.36 | 2505.90 | .001 | .92 |
| | Pipeline | 99349.05 | 1.72 | 358.46 | 57925.70 | 6385.48 | .001 | .97 |
| | Dataquality * Pipeline | 13224.60 | 1.22 | 255.05 | 10836.35 | 2094.07 | .001 | .91 |
| **L_Postcentral** | Dataquality | 10142.46 | 1 | 209 | 10142.46 | 2415.08 | .001 | .92 |
| | Pipeline | 156875.52 | 1.39 | 289.48 | 113260.61 | 5805.66 | .001 | .97 |
| | Dataquality * Pipeline | 26137.50 | 1.14 | 238.5 | 22900.77 | 2092.21 | .001 | .91 |
| **R_Postcentral** | Dataquality | 7852.87 | 1 | 209 | 7852.87 | 2482.27 | .001 | .92 |
| | Pipeline | 103496.80 | 1.73 | 360.512 | 60000.39 | 5897.63 | .001 | .97 |
| | Dataquality * Pipeline | 18480.19 | 1.22 | 254.96 | 15148.75 | 2160.06 | .001 | .91 |
| **L_Prefrontal** | Dataquality | 5896.11 | 1 | 209 | 5896.11 | 940.82 | .001 | .82 |
| | Pipeline | 304639.76 | 2.23 | 466.93 | 136358.99 | 7094.30 | .001 | .971 |
| | Dataquality * Pipeline | 22990.88 | 2.05 | 427.42 | 11242.03 | 1385.29 | .001 | .87 |
| **R_Prefrontal** | Dataquality | 9427.96 | 1 | 209 | 9427.96 | 1138.19 | .001 | .85 |
| | Pipeline | 320014.16 | 2.47 | 516.39 | 106671.39 | 6372.46 | .001 | .97 |
| | Dataquality * Pipeline | 27136.18 | 2.08 | 435.42 | 13025.30 | 1350.33 | .001 | .87 |
| **L_Parietal** | Dataquality | 12476.39 | 1 | 209 | 12476.39 | 2340.76 | .001 | .92 |
| | Pipeline | 191455.34 | 1.82 | 380.57 | 63818.45 | 4977.36 | .001 | .96 |
| | Dataquality * Pipeline | 36817.42 | 1.31 | 272.86 | 12272.47 | 2447.24 | .001 | .92 |
| **R_Parietal** | Dataquality | 11767.93 | 1 | 209 | 11767.93 | 1995.30 | .001 | .91 |
| | Pipeline | 128718.31 | 1.87 | 389.71 | 69031.52 | 4274.94 | .001 | .95 |
| | Dataquality * Pipeline | 29159.22 | 1.31 | 273.77 | 22260.76 | 1898.58 | .001 | .90 |
| **L_Temporal** | Dataquality | 14068.54 | 1 | 209 | 14068.54 | 2080.40 | .001 | .91 |
| | Pipeline | 198737.11 | 1.39 | 291.41 | 142533.85 | 5154.73 | .001 | .96 |
| | Dataquality * Pipeline | 46434.51 | 1.54 | 1.54 | 30139.17 | 2498.46 | .001 | .92 |
| **R_Temporal** | Dataquality | 15308.74 | 1 | 209 | 15308.74 | 2544.80 | .001 | .92 |
| | Pipeline | 134571.33 | 1.45 | 302.51 | 92972.12 | 5031.37 | .001 | .96 |
| | Dataquality * Pipeline | 35014.42 | 1.47 | 307.20 | 23821.45 | 2198.76 | .001 | .91 |
| **L_Occipital** | Dataquality | 11758.99 | 1 | 209 | 11758.99 | 1798.91 | .001 | .90 |
| | Pipeline | 111059.04 | 1.98 | 414.39 | 37019.68 | 3088.56 | .001 | .94 |
| | Dataquality * Pipeline | 20760.05 | 1.65 | 344.32 | 12601.23 | 1267.55 | .001 | .86 |
| **R_Occipital** | Dataquality | 14290.18 | 1 | 209 | 14290.18 | 1945.20 | .001 | .90 |
| | Pipeline | 115559.02 | 1.78 | 371.82 | 64955.41 | 3176.07 | .001 | .94 |
| | Dataquality * Pipeline | 20833.70 | 1.53 | 318.75 | 13660.28 | 1209.71 | .001 | .85 |

**Supplementary Table 4.** Two-way repeated measures ANOVA. The table summarizes main effects of within subject factors (data quality, pipeline) and their interaction (data quality * pipeline) on Streamline Density Index of each connectivity cluster. When sphericity was not satisfied (significant Mauchly’s Test) degrees of freedom were Greenhouse-Geisser corrected.

| DATA QUALITY * PIPELINES | | | | | | |
|---|---|---|---|---|---|---|
| **Cluster** | **Pipeline** | **df** | **error** | **F** | **Sig** | $\eta_p^2$ |
| **L_Paracentral** | CSD-THR | 1 | 209 | 161.98 | .001 | .89 |
| | CSD-WTA | 1 | 209 | 172.09 | .001 | .89 |
| | DTI-THR | 1 | 209 | 3.22 | .074 | .02 |
| | DTI-WTA | 1 | 209 | .58 | .447 | .00 |
| **R_Paracentral** | CSD-THR | 1 | 209 | 244.89 | .001 | .92 |
| | CSD-WTA | 1 | 209 | 117.92 | .001 | .85 |
| | DTI-THR | 1 | 209 | .000 | .984 | .00 |
| | DTI-WTA | 1 | 209 | 1.42 | .001 | .06 |
| **L_Postcentral** | CSD-THR | 1 | 209 | 237.05 | .001 | .92 |
| | CSD-WTA | 1 | 209 | 9.55 | .001 | .31 |
| | DTI-THR | 1 | 209 | 51 | .025 | .02 |
| | DTI-WTA | 1 | 209 | 1.31 | .001 | .06 |
| **R_Postcentral** | CSD-THR | 1 | 209 | 2542.90 | .001 | .92 |
| | CSD-WTA | 1 | 209 | 358.29 | .001 | .63 |
| | DTI-THR | 1 | 209 | 18.83 | .001 | .08 |
| | DTI-WTA | 1 | 209 | .04 | .85 | .00 |
| **L_Prefrontal** | CSD-THR | 1 | 209 | 2074.72 | .00 | .91 |
| | CSD-WTA | 1 | 209 | 962.80 | .00 | .82 |
| | DTI-THR | 1 | 209 | 146.79 | .00 | .41 |
| | DTI-WTA | 1 | 209 | 108.57 | .00 | .34 |
| **R_Prefrontal** | CSD-THR | 1 | 209 | 2117.36 | .00 | .91 |
| | CSD-WTA | 1 | 209 | 1184.61 | .00 | .85 |
| | DTI-THR | 1 | 209 | 316.09 | .00 | .60 |
| | DTI-WTA | 1 | 209 | 234.66 | .00 | .53 |
| **L_Parietal** | CSD-THR | 1 | 209 | 2911.23 | .00 | .93 |
| | CSD-WTA | 1 | 209 | 2.59 | .11 | .01 |
| | DTI-THR | 1 | 209 | .69 | .41 | .00 |
| | DTI-WTA | 1 | 209 | 8.31 | .00 | .04 |
| **R_Parietal** | CSD-THR | 1 | 209 | 2320.54 | .00 | .92 |
| | CSD-WTA | 1 | 209 | 16.12 | .00 | .07 |
| | DTI-THR | 1 | 209 | 37.50 | .00 | .15 |
| | DTI-WTA | 1 | 209 | 18.14 | .00 | .08 |
| **L_Temporal** | CSD-THR | 1 | 209 | 3143.10 | .00 | .94 |
| | CSD-WTA | 1 | 209 | 533.97 | .00 | .72 |
| | DTI-THR | 1 | 209 | 346.50 | .00 | .62 |
| | DTI-WTA | 1 | 209 | 11.64 | .00 | .05 |
| **R_Temporal** | CSD-THR | 1 | 209 | 2976.58 | .00 | .93 |
| | CSD-WTA | 1 | 209 | 5.11 | .02 | .02 |
| | DTI-THR | 1 | 209 | 356.96 | .00 | .63 |
| | DTI-WTA | 1 | 209 | 39.67 | .00 | .16 |
| **L_Occipital** | CSD-THR | 1 | 209 | 1990.83 | .00 | .90 |
| | CSD-WTA | 1 | 209 | 485.39 | .00 | .70 |
| | DTI-THR | 1 | 209 | 83.12 | .00 | .28 |
| | DTI-WTA | 1 | 209 | 2.69 | .10 | .01 |
| **R_Occipital** | CSD-THR | 1 | 209 | 1845.52 | .00 | .90 |
| | CSD-WTA | 1 | 209 | 977.03 | .00 | .82 |
| | DTI-THR | 1 | 209 | 208.74 | .00 | .50 |
| | DTI-WTA | 1 | 209 | 6.88 | .10 | .03 |

**Supplementary Table 5.** Simple main effects of data quality on Streamlines Density Index within each pipeline tested. These multivariate tests are based on pairwise comparisons linearly independent on estimated marginal means.

**PIPELINES * DATA QUALITY**

| Cluster | Data quality | df | error | F | sig | $\eta_p^2$ |
|---|---|---|---|---|---|---|
| **L_Paracentral** | High Quality | 3.00 | 207.00 | 1497.97 | .00 | .96 |
| | Downsampled | 3.00 | 207.00 | 2822.71 | .00 | .98 |
| **R_Paracentral** | High Quality | 3.00 | 207.00 | 1484.33 | .00 | .96 |
| | Downsampled | 3.00 | 207.00 | 3488.24 | .00 | .98 |
| **L_Postcentral** | High Quality | 3.00 | 207.00 | 1382.04 | .00 | .95 |
| | Downsampled | 3.00 | 207.00 | 2793.55 | .00 | .98 |
| **R_Postcentral** | High Quality | 3.00 | 207.00 | 802.96 | .00 | .92 |
| | Downsampled | 3.00 | 207.00 | 2821.12 | .00 | .98 |
| **L_Prefrontal** | High Quality | 3.00 | 207.00 | 4325.74 | .00 | .98 |
| | Downsampled | 3.00 | 207.00 | 5114.72 | .00 | .99 |
| **R_Prefrontal** | High Quality | 3.00 | 207.00 | 3834.40 | .00 | .98 |
| | Downsampled | 3.00 | 207.00 | 7090.49 | .00 | .99 |
| **L_Parietal** | High Quality | 3.00 | 207.00 | 1314.34 | .00 | .95 |
| | Downsampled | 3.00 | 207.00 | 2751.43 | .00 | .98 |
| **R_Parietal** | High Quality | 3.00 | 207.00 | 1616.22 | .00 | .96 |
| | Downsampled | 3.00 | 207.00 | 2894.70 | .00 | .98 |
| **L_Temporal** | High Quality | 3.00 | 207.00 | 4446.99 | .00 | .98 |
| | Downsampled | 3.00 | 207.00 | 4512.46 | .00 | .98 |
| **R_Temporal** | High Quality | 3.00 | 207.00 | 3910.58 | .00 | .98 |
| | Downsampled | 3.00 | 207.00 | 5663.54 | .00 | .99 |
| **L_Occipital** | High Quality | 3.00 | 207.00 | 2268.18 | .00 | .97 |
| | Downsampled | 3.00 | 207.00 | 2464.22 | .00 | .97 |
| **R_Occipital** | High Quality | 3.00 | 207.00 | 1398.45 | .00 | .95 |
| | Downsampled | 3.00 | 207.00 | 2562.40 | .00 | .97 |

**Supplementary Table 6.** Simple main effects of parcellation pipelines Streamlines Density Index within each kind of dataset tested. These multivariate tests are based on pairwise comparisons linearly independent on estimated marginal means.
